# Harnessing Inflammatory Monocytes to Overcome Resistance to Anti-PD-1 Immunotherapy

**DOI:** 10.64898/2026.02.05.704029

**Authors:** Matthew P Zimmerman, Amy Y Huang, Emily K Cox, Rose Al Abosy, Wan Lin Chong, Alexander G Bastian, Katherine Vietor, Yacine Choutri, Jenna Collier, Vasyl Zhabotynsky, Haiyang Wang, Megan Fung, Sarah A Weiss, Emily J Robitschek, Jia-Ren Lin, Tuulia Vallius, Shishir M Pant, Peter K Sorger, Willy Hugo, Debattama R Sen, W. Nicholas Haining, Arlene H Sharpe, Brian C Miller

**Affiliations:** Department of Cell Biology and Physiology, University of North Carolina at Chapel Hill, Chapel Hill, North Carolina, USA; Lineberger Comprehensive Cancer Center, University of North Carolina at Chapel Hill, Chapel Hill, North Carolina, USA; Department of Medical Oncology, Dana Farber Cancer Institute, Boston, MA; Harvard Ludwig Center and Laboratory of Systems Pharmacology, Harvard Medical School, Boston, MA; Department of Immunology, Blavatnik Institute, Harvard Medical School, Boston, MA, USA; Gene Lay Institute of Immunology and Inflammation, Brigham and Women’s Hospital, Massachusetts General Hospital and Harvard Medical School, Boston, MA, USA; Department of Medicine, Division of Oncology, University of North Carolina at Chapel Hill, Chapel Hill, North Carolina, USA; Department of Genetics, University of North Carolina at Chapel Hill, Chapel Hill, North Carolina, USA; Department of Microbiology and Immunology, University of North Carolina at Chapel Hill, Chapel Hill, North Carolina, USA; Department of Pediatric Oncology, Dana Farber Cancer Institute, Boston, MA, USA; Department of Medicine, UCLA David Geffen School of Medicine, Los Angeles, California, USA; Krantz Family Center for Cancer Research, Department of Medicine, Massachusetts General Hospital Boston, MA, USA; Department of Obstetrics and Gynecology, University of Washington, Seattle, WA, USA; Carbone Cancer Center, University of Wisconsin at Madison, Madison, WI and Cancer Biology Graduate Program, University of Wisconsin at Madison, Madison, WI, USA; Department of Systems Biology, Harvard Medical School, Boston, MA, USA; Arsenal Biosciences, San Francisco, CA, USA; Department of Medicine, University of California San Francisco, San Francisco, CA, USA; Department of Paediatrics, University of Toronto, Toronto, Ontario, Canada; Genentech, San Francisco, CA, USA; Reglab, Stanford University, Stanford, CA, USA; GC Therapeutics, Cambridge, MA, USA; Massachusetts Institute of Technology, Cambridge, MA, USA; Department of Pathology, Brigham and Women’s Hospital, Boston, MA, USA; Broad Institute of Harvard and Massachusetts Institute of Technology, Cambridge, MA, USA; Division of Medical Sciences, Harvard Medical School, Boston, MA, USA; Division of Population Sciences, Dana-Farber Cancer Institute, Boston, MA, USA; Department of Medicine, Harvard Medical School, Boston, MA, USA; Center for Immunology and Inflammatory Diseases, Massachusetts General Hospital, Boston, MA, USA

**Author notes:** Corresponding Author: Brian C Miller.

**Keywords:** Immune Checkpoint Inhibitor, Monocytes, T-Lymphocytes, Major Histocompatibility Complex - MHC

## Abstract

**Background:** Resistance to immune checkpoint inhibitors represents a major therapeutic challenge, as less than 50% of patients with melanoma achieve long-term response to immune checkpoint inhibitor therapy. One mechanism of acquired resistance involves somatic mutations, such as loss of beta-2 microglobulin (*B2m*), that enable tumor cells to evade T cell-mediated killing.

**Methods:** This study used single-cell RNA-seq, flow cytometry, and *ex vivo* functional assays to characterize tumor-infiltrating immune cells in antigen presentation-deficient tumors. Tumor-bearing mice were treated with anti-PD-1 or CD40 agonist antibodies and cell depletion or cytokine blocking antibodies to define mechanisms of action. Analysis of published human RNA-seq datasets was performed to dissect the contributions of inflammatory monocytes to patient outcomes.

**Results:** We found an increase in immunosuppressive macrophages in *B2m*-null tumors. We hypothesized that repolarizing myeloid cells may restore control of tumor growth. Treatment with CD40 agonist antibody, which promotes differentiation of monocytes and macrophages towards a proinflammatory phenotype, reduced tumor growth and improved survival in *B2m*-null melanoma and colorectal cancer models. Unexpectedly, both CD8^+^ T cells and NK cells, but not CD4^+^ T cells, were required for the efficacy of CD40 agonist, even though CD8^+^ T cells cannot directly recognize antigen presentation-deficient tumor cells. Instead, these lymphocytes control tumor growth via secretion of IFNγ, as depletion of IFNγ inhibited the therapeutic effect of CD40 agonist. IFNγ receptor (*Ifngr1*) expression was required on host cells, not tumor cells, for CD40 agonist-mediated tumor control. Single-cell analysis identified a distinct population of inflammatory monocytes that were enriched for an IFNγ response signature in CD40 agonist-treated tumors, suggesting that these cells may be important for tumor control. Analysis of human bulk and single-cell RNA-seq datasets demonstrated that an inflammatory monocyte signature derived from our data was associated with improved patient outcomes and response to immune checkpoint inhibitors.

**Conclusions:** These data demonstrate that CD8^+^ T cells contribute to tumor control even in the absence of direct antigen presentation by tumor cells. More broadly, our work suggests that strategies to activate the effector functions of inflammatory monocytes may limit tumor growth and overcome acquired resistance to immune checkpoint inhibitors.

## BACKGROUND

Resistance to immune checkpoint inhibitors (ICI) remains a major therapeutic challenge in oncology. Tumor cells can acquire resistance to ICI by preventing presentation of tumor antigens to CD8^+^ T cells, the predominant cytotoxic effectors of the anti-tumor immune response. Loss of antigen presentation can result from decreased expression of Major Histocompatibility Complex I (MHC-I) due to mutations in the essential subunit *B2m*, epigenetic silencing, or heterozygous loss of MHC-I alleles.[2–8] Consistent with this, loss of cell surface expression of MHC-I has been reported in 40-90% of human cancers.[9] Prior work has demonstrated that NK cells, γδ T cells, CD8^+^ T cells, or CD4^+^ T cells can kill MHC-I-deficient tumor cells.[5,10–13] However, therapeutic approaches that effectively target these cells have not yet been translated into the clinic.

CD40 is a Tumor Necrosis Factor (TNF) receptor superfamily member expressed on the surface of many immune cells, including macrophages, monocytes, dendritic cells, and B cells. Engagement by its cognate ligand, CD40L, activates CD40 signaling that results in maturation and proinflammatory differentiation of immune cells. CD40 agonists are currently being explored in clinical trials for treatment of ICI refractory patients, including anti-PD-1 resistant melanoma and pancreatic cancer. However, therapeutic responses in these clinical studies have been limited, likely due to dose-limiting toxicities.[14–17] Newly-developed CD40 agonists designed to improve this therapeutic window are in early phase clinical trials.[18] In preclinical models, CD40 agonists control tumor growth through both CD8^+^ T cell-dependent and -independent mechanisms.[11,19–23] Defining the mechanisms by which CD40 agonists mediate tumor control is therefore critical to improving their clinical efficacy.

There is growing interest in the role of inflammatory monocytes to directly control cancer growth. Preclinical studies demonstrate that interferon-stimulated monocytes can kill a variety of human cancer cell lines *in vitro* and *in vivo*.[24–26] Consistent with these findings, a recent phase I clinical trial of autologous interferon-activated monocytes administered intraperitoneally to ovarian cancer patients suggests that this cellular therapy can attenuate disease progression in a subset of patients.[27] Proposed mechanisms of action for the cytotoxic effects of monocytes include direct induction of cancer cell apoptosis by TRAIL (TNF-Related Apoptosis-Inducing Ligand), generation of nitric oxide, cross-presentation of antigens to CD8^+^ T cells, and degradation of extracellular matrix networks to enhance chemotherapy efficacy.[27–30] Consequently, cancer cells impair monocyte activation to counter these anti-tumor functions.[30]

To identify strategies to overcome resistance due to loss of antigen presentation, we generated *B2m*-null murine cancer cell lines and characterized the immune microenvironment of antigen presentation-deficient tumors. We identified a robust infiltration of immunosuppressive macrophages in *B2m*-null melanoma tumors. To therapeutically reprogram these immunosuppressive macrophages and restore tumor control, we treated *B2m*-null tumor-bearing mice with CD40 agonist. We found that CD40 agonist monotherapy was sufficient to slow, but not eradicate, *B2m*-null tumor growth. Surprisingly, CD8^+^ T cells were still required for therapeutic efficacy, despite their inability to directly recognize tumor cells lacking MHC-I. This effect required IFN𝛄 signaling in host immune cells but not tumor cells. CD40 agonist treatment induced an influx of interferon-responsive inflammatory monocytes into the tumor microenvironment, and we propose that these cells contribute to tumor control. Consistent with this model, cancer patients with higher inflammatory monocyte signatures had a better prognosis across many different tumor types, underscoring the clinical relevance of these cells.

## RESULTS

### Loss of *B2m* by tumor cells increases infiltration by immunosuppressive macrophages

To model the loss of antigen presentation, we deleted *B2m* in B16F10 melanoma tumor cells using CRISPR-Cas9 (Supp. Fig. 1A). Consistent with prior findings in melanoma tumor models and patients,[4,5,10,31] *B2m*-null tumors did not respond to anti-PD-1 immunotherapy (combined with GVAX vaccination), unlike control B16 tumors (Fig. 1A, 1B). To determine how loss of *B2m* changes the tumor immune microenvironment, we performed single-cell RNA-seq on tumor-infiltrating immune cells (Fig. 1C, Supp. Fig. 1B-D). We hypothesized that in the absence of a productive anti-tumor immune response, *B2m*-null tumors would contain more pro-tumor immune cells, including immunosuppressive tumor-associated macrophages (TAMs). Indeed, differential expression analysis of total myeloid cells from control versus *B2m*-null tumors revealed increased expression of canonical genes associated with immunosuppressive TAMs (e.g. *Maf*, *Cd163*, *Mrc1*) in *B2m*-null tumors, whereas control tumors were enriched for proinflammatory macrophage genes (e.g. *Cxcl10*, *Irf8*, *H2-D1*) (Fig. 1D). In addition, a gene signature of classically-activated macrophages (IFNγ+LPS stimulated) was enriched in TAMs from control tumors while a signature of alternatively activated (IL-4 stimulated) macrophages was enriched in TAMs from *B2m*-null tumors (Fig. 1E). Flow cytometry confirmed an increased proportion of CD206+ and MHC-II-TAMs in *B2m*-null tumors (Fig. 1F, 1G, Supp. Fig. 1E). These data demonstrate that TAMs in *B2m*-null tumors exhibit a more immunosuppressive phenotype than TAMs found in control tumors.

**Figure 1.**
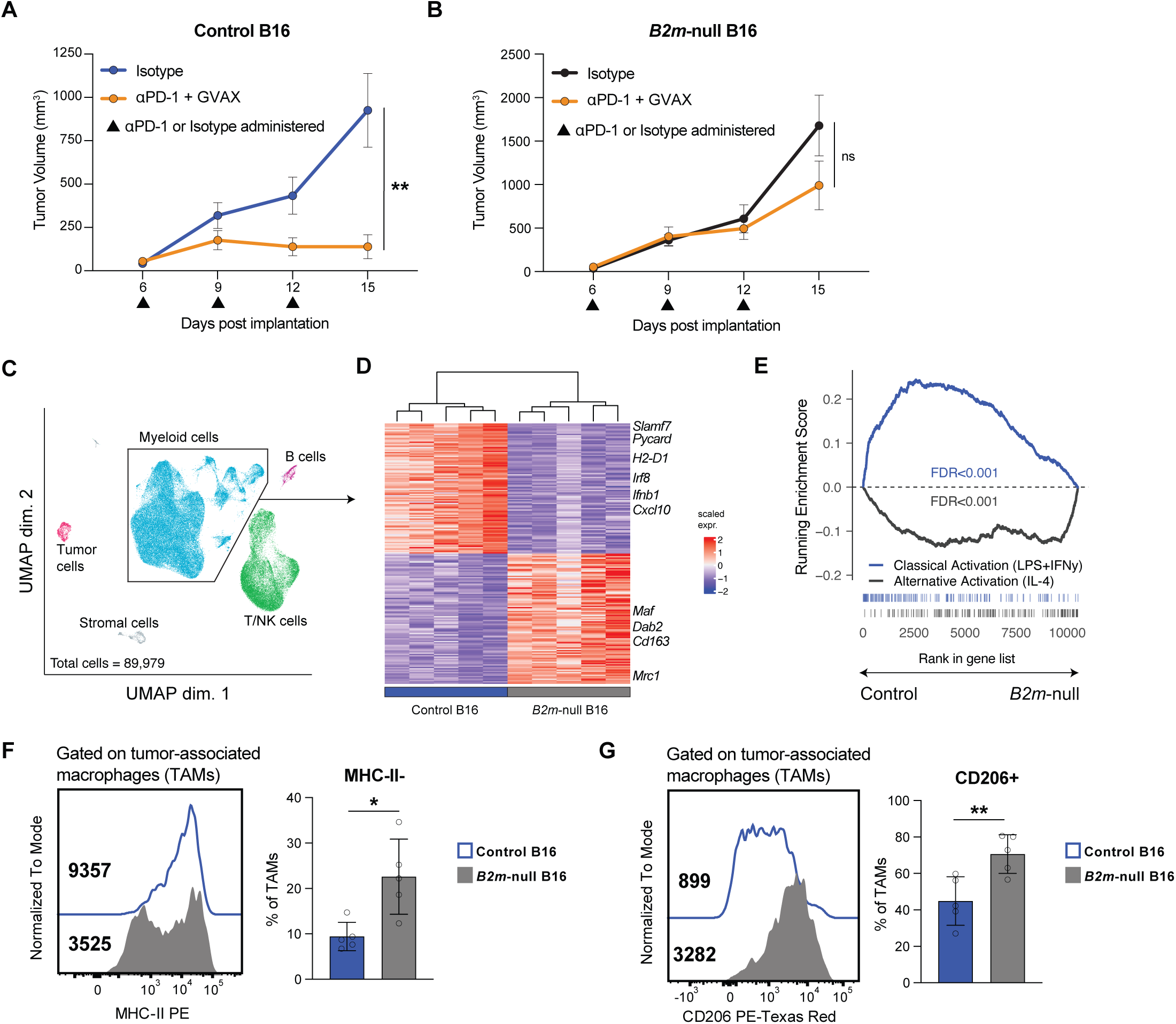
MHC-I-deficient tumors are enriched in immunosuppressive macrophages. **(A, B)** Growth curves of control **(A)** or *B2m*-null **(B)** B16 tumors implanted into C57BL/6J mice and treated with either anti-PD-1 (αPD-1) or isotype control antibodies days 6, 9 and 12 post-implantation. αPD-1 treated mice were also vaccinated with GVAX on days 1 and 4 post-implantation. Representative results from one of three independent experiments. **(C)** UMAP of single-cell RNA-sequencing profiles from 89,979 cells isolated from day 14 isotype- or CD40 agonist-treated control (n = 5 isotype, 2 = CD40 agonist) and *B2m*-null (n = 5 isotype, 4 = CD40 agonist) B16 melanoma tumor samples. Each point represents an individual cell spatially-clustered and color-coded according to cell type annotation across all samples. **(D)** Heatmap of top differentially expressed genes in the myeloid cell cluster in isotype-treated control (n = 5) or *B2m*-null (n = 5) B16 tumors. **(E)** Gene set enrichment analysis of classical and alternative activation macrophage signatures (GSE69607) in the ranked list of genes differentially expressed in pseudobulk data from myeloid cells in control versus *B2m*-null B16 tumors. **(F,G)** Flow cytometry analysis of MHC-II **(F)** and CD206 **(G)** expression in control or *B2m*-null B16 TAMs. Representative flow plots (left) and summary (right) from one of two independent experiments. * p < 0.05; ** p < 0.01; n.s. not significant.

### *B2m*-null tumor growth is controlled by CD40 agonist treatment

As TAMs in *B2m*-null tumors are skewed towards an immunosuppressive phenotype, we hypothesized that repolarizing myeloid cells towards a proinflammatory state could improve tumor control. To test this, we treated *B2m*-null tumor-bearing mice with CD40 agonist.[19,21,32,33] In both *B2m*-null B16 and MC38 tumors, CD40 agonist suppressed tumor growth and prolonged survival (Fig. 2A, B). Given that *B2m* is also required for the expression of many non-classical MHC molecules, which can stimulate or suppress anti-tumor effector cells,[34,35] we also generated dual-knockout B16 cells lacking both *H2-K1* and *H2-D1*, the classical MHC-I alleles expressed in C57BL/6J mice (Supp. Fig. 2A). These tumors were also controlled by CD40 agonist treatment, indicating that the observed effect is unlikely to be mediated by loss of non-classical MHC-I in *B2m*-null tumor cells (Supp. Fig. 2B).

**Figure 2.**
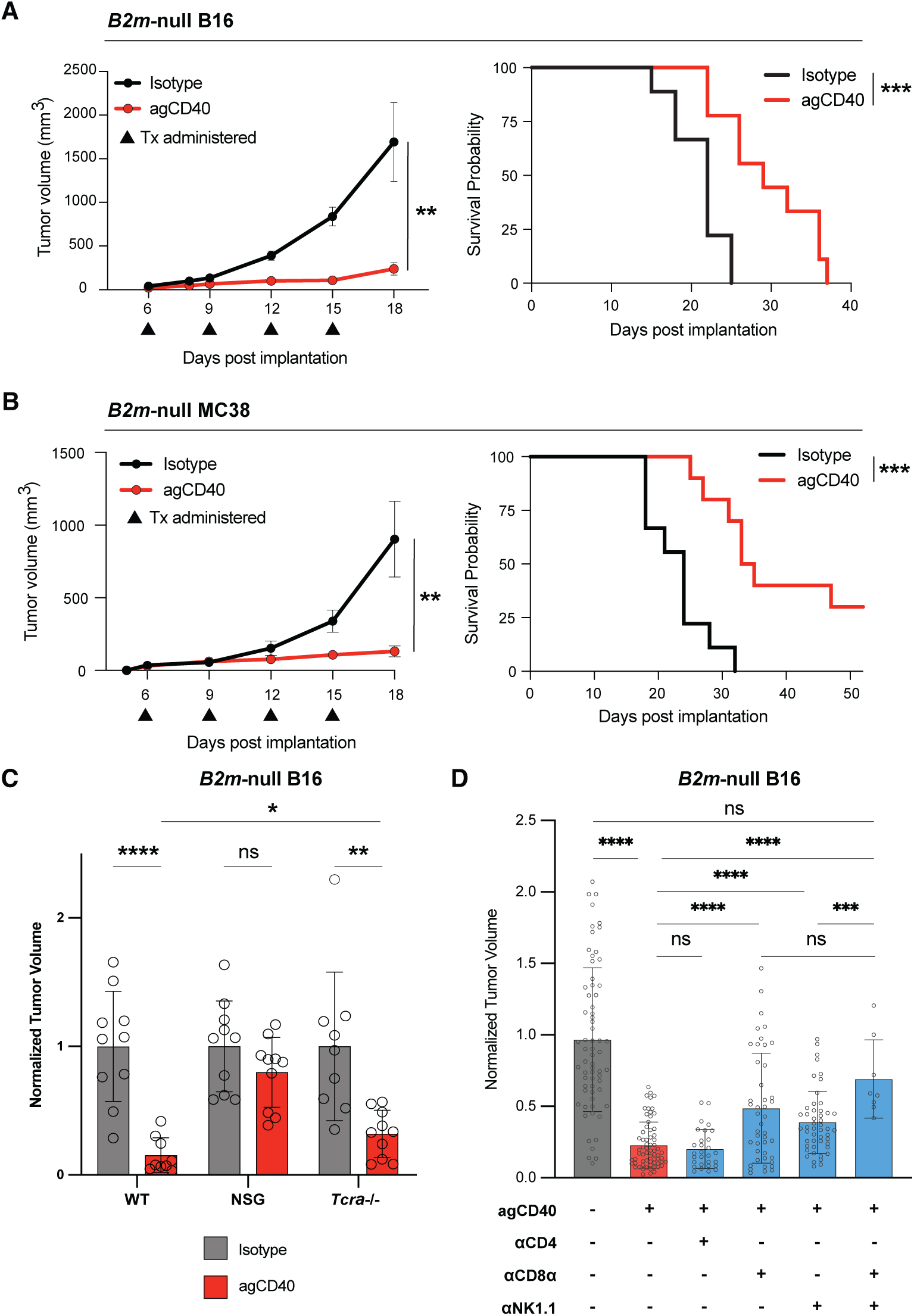
CD40 agonist monotherapy controls MHCI-deficient tumor growth and requires CD8^+^ T cells and NK cells for efficacy. **(A)** Growth curve (left; n = 5 isotype-, 5 CD40 agonist (agCD40)-treated) and Kaplan-Meier survival analysis (right; n = 9 isotype-, 9 agCD40-treated) of wild-type C57BL/6J mice bearing subcutaneous *B2m*-null B16 tumors treated with CD40 agonist (agCD40) or isotype control antibodies. Representative results from one of at least three independent experiments for tumor volume analyses and one of two independent experiments for survival studies. **(B)** Growth curve (left) and Kaplan-Meier survival analysis (right) of wild-type C57BL/6J mice bearing subcutaneous *B2m*-null MC38 tumors treated with agCD40 (n = 10) or isotype control (n = 9) antibodies. Representative results from one of two independent experiments. **(C)** *B2m*-null B16 harvest tumor volumes from wild-type C57BL/6J (n = 10 isotype-, 9 agCD40-treated), NSG (NOD.Cg-Prkdc^scid^ Il2rg^tm1Wjl^/SzJ; n = 10 isotype-, 10 agCD40-treated), or *Tcra*^-/-^(B6.129S2-Tcra^tm1Mom^/J; n = 9 isotype-, 10 agCD40-treated) mice treated with isotype control or CD40 agonist antibody. Tumor volumes are normalized to the mean volume of isotype-treated mice within respective mouse genotype. Results compiled from three independent experiments. **(D)** *B2m*-null B16 tumor volumes from wild-type C57BL/6J mice treated with isotype control or CD40 agonist antibody in conjunction with anti-CD8α (αCD8α), anti-CD4 (αCD4), and/or anti-NK1.1 (αNK1.1) depleting antibodies. Tumor volumes are normalized to the mean volume of isotype-treated mice for each experiment. Results pooled from eight independent experiments. * p < 0.05; ** p < 0.01; *** p < 0.001; **** p < 0.0001; n.s. not significant.

### T cells and NK cells are required for CD40 agonist efficacy

Because CD40 agonists can act via both T cell-dependent and -independent mechanisms,[11,19–23,36] we next asked which cells are required for CD40 agonist-mediated control of *B2m*-null tumors. We first implanted *B2m*-null B16 tumors into control C57BL/6J, NSG mice (which lack functional T cells, B cells, and Natural Killer cells), or *Tcra*^-/-^ mice, which lack conventional CD4^+^ and CD8^+^ T cells (Fig. 2C, Supp Fig. 3A). The efficacy of CD40 agonist was completely lost in NSG mice, indicating NK cells and/or adaptive immune cells are required for tumor control. *Tcra*^-/-^ mice that lack conventional ɑβ T cells still responded to CD40 agonist treatment, although with a significant increase in tumor size compared with CD40-treated control mice. These data suggest that both NK cells and T cells may contribute to CD40 agonist-mediated control of *B2m*-null tumors.

To further define the specific cell types required, we next performed antibody depletion studies. Depletion of CD4^+^ T cells, B cells, or γδ T cells did not alter the therapeutic effect of CD40 agonist treatment. In contrast, depleting either CD8^+^ T cells or NK cells partially impaired tumor control by CD40 agonist, whereas depleting both cell types resulted in larger tumors (Fig. 2D, Supp. Fig. 3B-G). These data suggest that both CD8^+^ T cells and NK cells are required for maximal efficacy of CD40 agonist. Supporting this hypothesis, in some experiments only the combination of CD8^+^ T cell and NK cell depletion abrogated CD40 agonist efficacy, while single depletion had no effect (Supp. Fig. 3B). Prior studies have shown that tumor clearance can require either one or both cell types, depending on tumor inoculum and initial tumor burden at treatment onset.[37] While NK cells have a well-described role to eliminate tumor cells lacking MHC-I expression, recent work suggests that CD8^+^ T cells can kill tumor cells via NKG2D-mediated cytotoxicity.[12,38] We observed that <15% of CD8^+^ T cells in our model express NKG2D, and expression did not increase with CD40 agonist treatment (Supp. Fig. 4A). These data suggest that CD40 agonist requires NK and CD8^+^ T cells to control tumor growth through an alternative mechanism.

### CD40 agonist treatment drives T cell infiltration into *B2m*-null tumors

To investigate the surprising role of CD8^+^ T cells to control tumors lacking antigen presentation, we performed single-cell RNA-seq on tumor-infiltrating immune cells from control and *B2m*-null tumors after isotype or CD40 agonist treatment (Fig. 3A). CD40 agonist treatment induced a dramatic remodeling of the tumor microenvironment in both control and *B2m*-null tumors (Fig. 3B). While *B2m*-null tumors have very low baseline T cell infiltration, CD40 agonist treatment led to a marked increase in T cells within these tumors (Fig. 3B, 3C). We confirmed the increased infiltration of T cells into the tumor microenvironment (TME) by flow cytometry. Both CD8^+^ T cells and CD4^+^ T cells increased as a percentage of total tumor-infiltrating immune cells, as well as in total numbers of cells normalized to tumor weight (Fig. 3D, Supp. Fig. 4B). In contrast to T cells, NK cell numbers in the TME did not change after CD40 agonist treatment (Fig. 3E).

**Figure 3.**
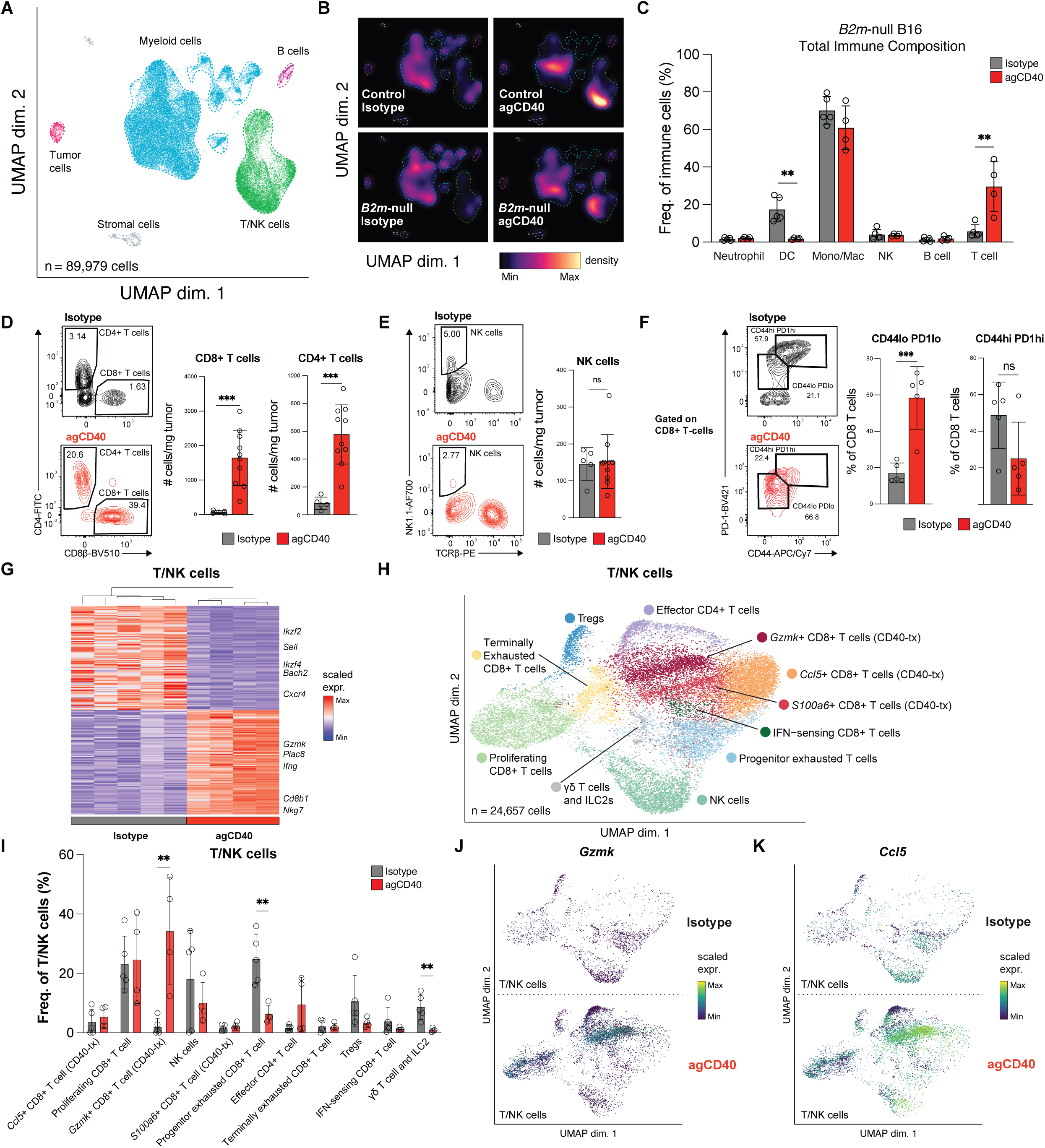
CD40 agonist induces an expansion of transcriptionally distinct, activated CD8^+^ T cells. **(A, B)** scRNA-seq profiles from 89,979 cells extracted from control and *B2m*-null B16 tumors treated with CD40 agonist (agCD40) or isotype control antibody. UMAP of major cell clusters (Tumor, Myeloid, B, T/NK, and Stromal cells) **(A)** and cell density via galaxy plot in UMAP space **(B)** (n = 5 control-iso-type, 2 control-agCD40, 5 *B2m*-null-isotype, and 4 *B2m*-null-agCD40). **(C)** Quantification of each major cell cluster from scRNAseq analysis as a frequency of total immune cells in *B2m-null* B16 tumors. **(D, E)** Quantification of normalized cell numbers for CD8^+^ T cells, CD4^+^ T cells **(D)** and NK cells **(E)** from *B2m*-null B16 tumors treated with agCD40 (n = 10) or isotype control (n = 5) antibody. Represen-tative flow plots reporting the percentage of each population as a frequency of all Live CD45+ cells (left) and summary of results (right) from one of at least three independent experiments. **(F)** CD44 and PD-1 expression on CD8^+^ TILs from *B2m*-null B16 tumors treated with agCD40 or isotype control antibody. Representative flow plot (left) and summary (right) from one of two independent experiments. **(G)** Heatmap of top differentially expressed genes from the T/NK cell cluster in isotype or agCD40 treated *B2m*-null tumors. **(H)** UMAP of scRNA-seq profiles of 24,657 cells reclustered from the T/NK cell subpopulation. **(I)** Quantification of each T/NK cell cluster as a frequen-cy of total T/NK cells. **(J, K)** Expression of *Gzmk* **(J)** and *Ccl5* **(K)** in individual cells from **(H)**.** p < 0.01; *** p < 0.001; n.s. not significant.

We next examined the activation states of lymphocytes after CD40 agonist treatment. Flow cytometry revealed a significant increase in CD44^lo^PD-1^lo^ CD8^+^ T cells in tumors and a decrease in TIM-3 expression after CD40 agonist treatment, consistent with decreased activation and/or exhaustion (Fig. 3F, Supp. Fig. 4C). However, transcripts associated with cytotoxic effector CD8^+^ T cells were upregulated in *B2m*-null tumors treated with CD40 agonist (e.g. *Ifng, Nkg7, Gzmk*), whereas transcripts found in quiescent lymphocytes were expressed in isotype-treated *B2m*-null tumors (e.g. *Ikzf2, Sell, Bach2, Ikzf4, Cxcr4*) (Fig. 3G). Reprojection and analysis of the T/NK cell clusters revealed a transcriptionally distinct population of CD8^+^ T cells in CD40 agonist-treated *B2m*-null tumors (Fig. 3H, 3I, Supp. Fig. 4D-F). Importantly, these cells did not fall into conventional progenitor or terminally exhausted cell states, as previously described in B16 tumors.[39] These CD8^+^ T cells expressed high levels of *Ccl5* and *Gzmk* (Fig. 3J, 3K). CCL5 has previously been shown to be upregulated by myeloid and CD8^+^ T cells in response to CD40 agonist treatment and is important for CD4^+^ T cell recruitment, while Granzyme K marks activated CD8^+^ T cells that are major cytokine producers but have reduced cytotoxicity.[23,40] Together, these results demonstrate that CD40 agonist not only expands the intratumoral T cell compartment, but also drives CD8^+^ T cells into a distinct activation state.

### IFNγ signaling on host cells is required for the efficacy of CD40 agonist

In addition to *Ccl5* and *Gzmk*, we noticed that *Ifng* (interferon gamma, IFNγ) was significantly upregulated across multiple lymphocyte clusters after CD40 agonist treatment (Fig. 3G, Fig. 4A). IFNγ can exert both direct cytostatic and cytotoxic effects on tumor cells as well as activate other immune cells.[41–44] To validate increased IFNγ within the TME *in vivo*, we scored all cells in our scRNAseq dataset using an IFNγ response signature. CD40 agonist treatment significantly increased the IFNγ response score both across all cell types as well as within myeloid and T/NK cells, consistent with increased IFNγ in the TME (Fig. 4B, 4C, Supp. Fig. 5A). *Ex vivo* restimulation of tumor-infiltrating lymphocytes with PMA and ionomycin confirmed that a greater percentage of CD4^+^ and CD8^+^ T cells extracted from CD40 agonist-treated tumors produced IFNγ relative to cells isolated from isotype-treated tumors. In contrast to our single-cell transcriptional data, however, we did not observe an increase in IFNγ production by NK cells (Fig. 4D).

**Figure 4.**
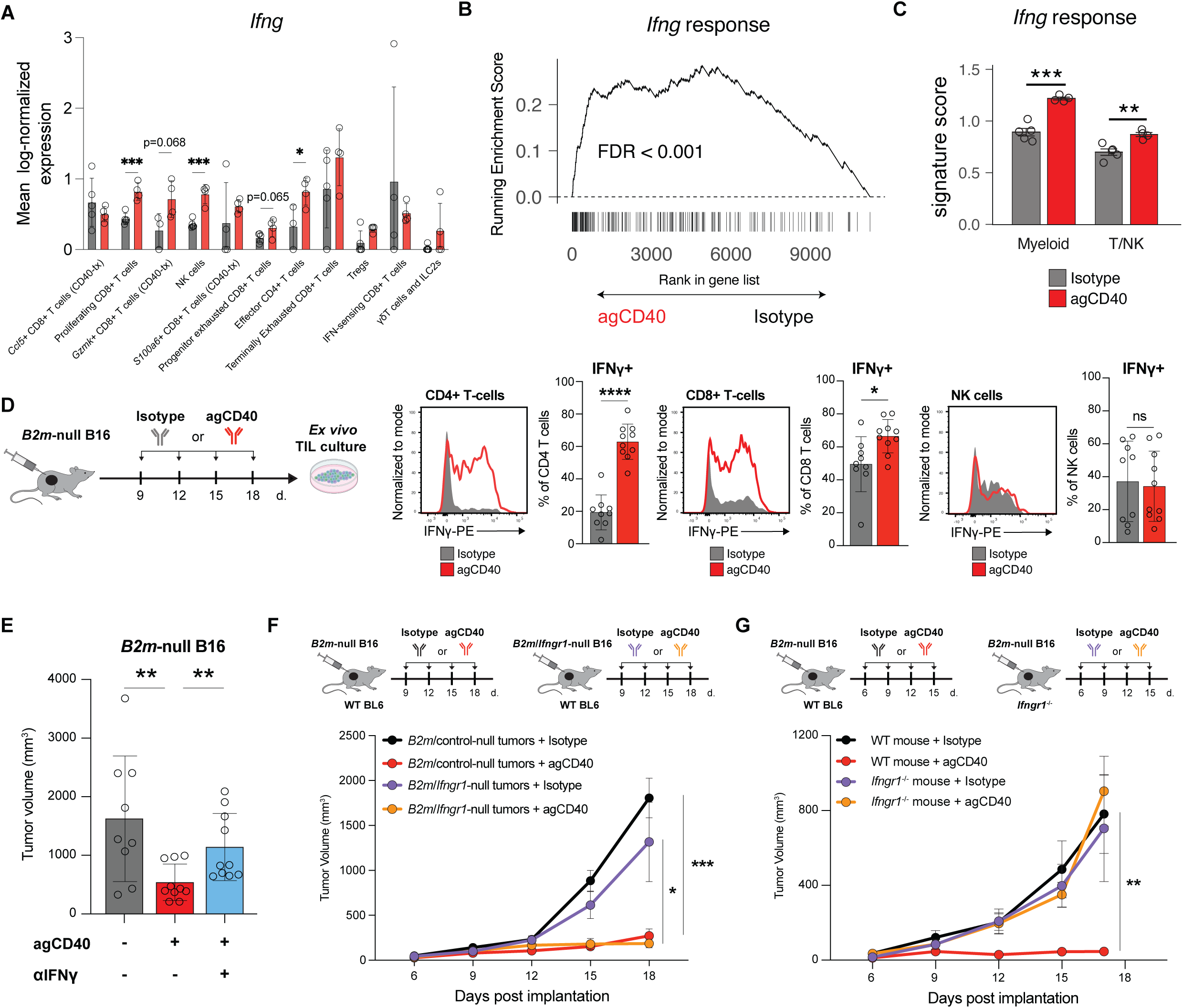
CD40 agonist-mediated control of MHC-I-deficient tumors requires host IFNγ signaling. **(A)** Expression of *Ifng* by different populations of T/NK cells from scRNAseq analysis. **(B)** GSEA of the Hallmark IFNγ response signature in the ranked list of genes differentially expressed in pseudobulk data from all cells isolated from agCD40 vs isotype treated *B2m*-null B16 tumors. **(C)** Per sample analysis of the Hallmark IFNγ response signature in myeloid cells or T/NK cells. **(D)** Frequency of IFNγ-producing CD4^+^ T-cells (left), CD8^+^ T-cells (center) and NK cells (right) extracted from isotype control- (n = 9) or agCD40-treated (n = 10) *B2m*-null B16 tumors and stimulated *ex vivo* for 6h with PMA/ionomy-cin. Representative flow plots and quantification of pooled results from two independent experiments shown. **(E)** Tumor volumes of *B2m*-null B16 tumors treated with agCD40 or isotype control antibody and IFNγ blocking antibody. Represen-tative day 12 tumor volumes from one of two independent experiments. **(F)** Growth curves of *B2m*-null or *B2m*/*Ifngr1*-null dual knockout B16 tumors treated with isotype control or agCD40. Representative results from one of two independent experiments. **(G)** Growth curves of *B2m*-null B16 tumors implanted into WT C57BL6/J or *Ifngr1*^-/-^ mice and treated with isotype control or agCD40. Representative results from one of two independent experiments. * p < 0.05; ** p < 0.01; *** p < 0.001; n.s. not significant.

To determine if IFNγ is required for CD40 agonist-mediated control of *B2m*-null tumors, we treated tumor-bearing mice with an IFNγ blocking antibody concurrently with CD40 agonist. IFNγ blockade completely abrogated the therapeutic effect of CD40 agonist (Fig. 4E). We next asked in which cellular compartment IFNγ signaling was required for CD40 agonist efficacy. We first generated dual-knockout *B2m*-null/*Ifngr1*-null B16 cells lacking a subunit of the IFNγ receptor necessary for downstream signaling (Supp. Fig. 5B). When these cells were implanted into wild-type mice, the tumors still responded to agonist CD40 treatment, demonstrating that the critical action of IFNγ is not directly on tumor cells (Fig. 4F). To test whether IFNγ signaling is required on host cells to mediate tumor control, we implanted *B2m*-null B16 tumors into *Ifngr1*^-/-^ mice and treated them with CD40 agonist. *B2m*-null tumors no longer responded to CD40 agonist when implanted in *Ifngr1*^-/-^ mice (Fig. 4G). Together, these data demonstrate that CD40 agonist treatment requires IFNγ signaling in host cells rather than tumor cells to control *B2m*-null tumor growth.

### CD40 agonist treatment promotes tumor infiltration by inflammatory monocytes

To determine which cells may mediate the effects of IFNγ, we examined the IFNγ response signature across cell clusters in our single-cell dataset and found the strongest enrichment in a cluster of tumor-infiltrating monocytes. This prompted a focused analysis of tumor-infiltrating myeloid cells (Fig. 5A-B, Supp. Fig. 6A-C). Across all myeloid cells, CD40 agonist treatment increased expression of interferon-activated proinflammatory monocyte and macrophage transcripts, including chemokines important for immune cell recruitment (*Cxcl16*), antigen presentation (*B2m, H2-Q7, Ifi30*) and markers of proinflammatory metabolism (*Nos2, Acod1*). Conversely, myeloid cells from isotype-treated tumors were enriched for markers of tolerogenic myeloid cells and plasmacytoid dendritic cells (*Siglech, Tcf4, Flt3*) (Fig. 5C).

**Figure 5.**
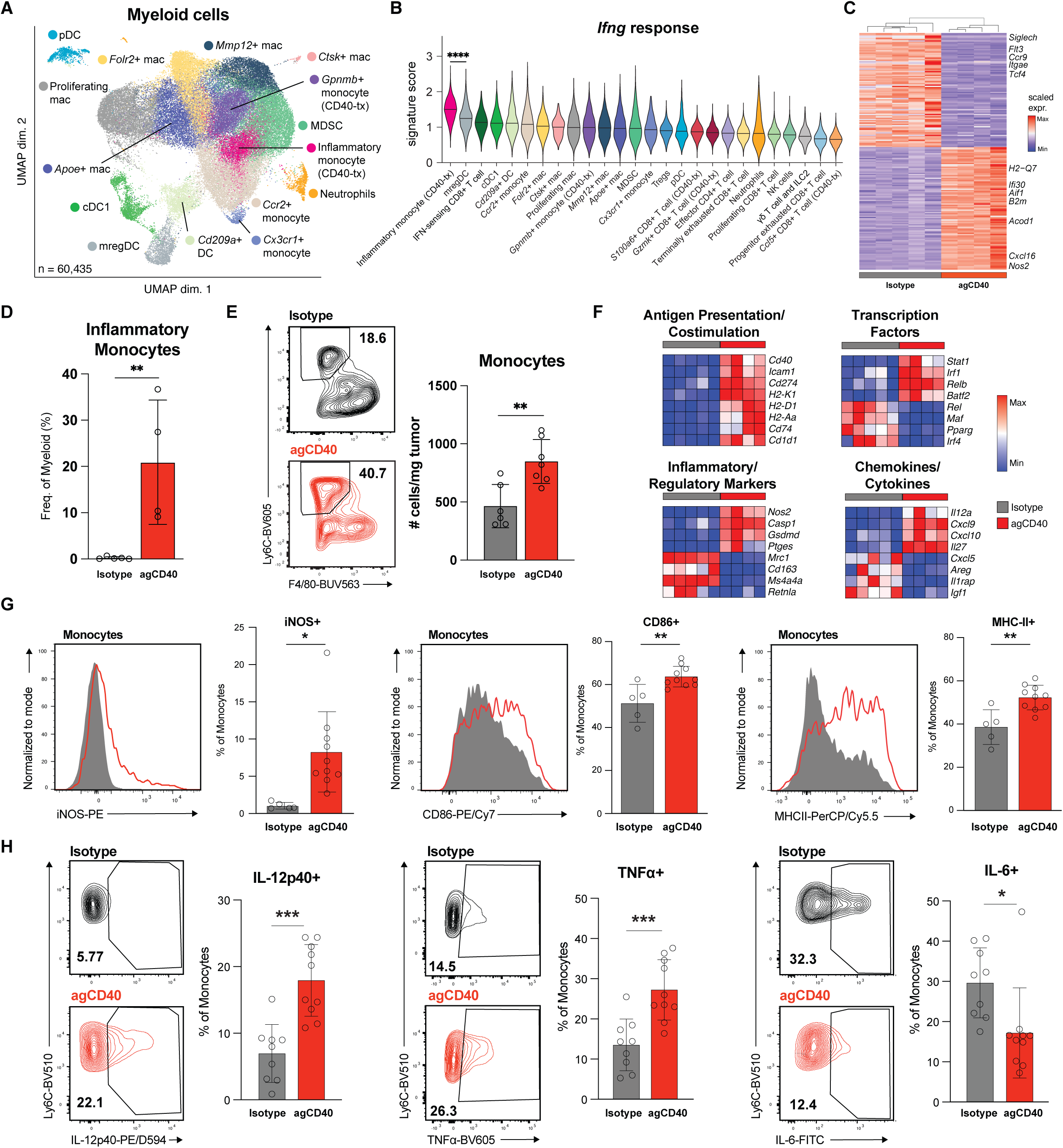
CD40 agonist drives expansion of inflammatory monocytes in MHC-I-deficient tumors. **(A)** UMAP of scRNA-seq profiles of 60,435 myeloid cells reclustered from the myeloid cell subpopulation. **(B)** Violin plot depicting the Hallmark *Ifng* response score in myeloid and T/NK cell subclusters. **(C)** Heatmap of top differentially-expressed genes in total myeloid cells between isotype- and agCD40-treated *B2m*-null B16 tumor samples from scRNAseq analysis. **(D)** Quantification of the inflammatory monocyte cell cluster from scRNAseq analysis of *B2m*-null B16 tumors as the frequency of total myeloid cells. **(E)** Number of tumor-infiltrating monocytes from isotype control- (n = 6) or agCD40-treated (n = 7) *B2m*-null B16 tumors normalized by individual tumor weight. Representative flow plots displaying percentage of inflammatory monocytes as a frequency of Live CD45+ cells (left) and summary quantifiation (right) from one of at least three independent experiments. **(F)** Heatmap illustrating the average transcript expression of the indicated genes across all monocyte subclusters per sample. Rows represent averaged z-scores. All indicated genes have a significant differential expression by DESeq2 (q<0.05) **(G)** Frequency of iNOS+ (left), CD86+ (center), and MHC-II+ (right) tumor-infiltrating monocytes from isotype control- or agCD40-treated *B2m*-null B16 tumors. Representative contour flow plots (left) and quantification of flow cytometry analysis (right) from one of two independent experiments. **(H)** Frequency of IL-12p40+ (left), TNFα+ (center), and IL-6+ (right) monocytes extracted from isotype control- (n = 9) or agCD40-treated (n = 10) *B2m*-null B16 tumors and stimulated *ex vivo* for 18h with LPS. Representative contour flow plots (left) and quantification of flow cytometry analysis (right) pooled between two independent experiments. * p < 0.05; ** p < 0.01; *** p < 0.001; n.s. not significant.

Further dissecting into myeloid cell subsets, we found that CD40 agonist treatment increased tumor infiltration by inflammatory monocytes, which represent approximately 20% of all myeloid cells in treated *B2m*-null tumors (Fig. 5D, 5E, Supp. Fig. 6D). Across all monocytes, CD40 agonist induced high levels of inflammatory markers and cytokines/chemokines (*Nos2, Cxcl9*, *Il12a)*, antigen presentation genes (*H2-Ab1*, *H2-Eb1*), and proinflammatory transcription factors (*Irf1, Stat1*), while downregulating regulatory markers and transcription factors (*Mrc1, Cd163, Pparg, Mafb*), consistent with immune activating functions of inflammatory monocytes (Fig. 5F). Flow cytometry confirmed repolarization of tumor-infiltrating myeloid cells, demonstrating increased expression of proinflammatory markers (iNOS, CD86, MHC-II) in monocytes and TAMs (Fig. 5G, Supp. Fig. 6E). Together, these results demonstrate that CD40 agonist treatment repolarizes myeloid cells towards a proinflammatory state, with a large influx of inflammatory monocytes.

To confirm that these monocytes are functionally proinflammatory, we sorted monocytes from isotype and CD40 agonist-treated *B2m*-null B16 tumors and cultured them *ex vivo*, with or without LPS restimulation. Monocytes isolated from CD40 agonist-treated tumors produced higher levels of the immunostimulatory cytokines IL-12 and TNFɑ and reduced IL-6 production, which can promote tumor growth, compared with monocytes from isotype control-treated tumors (Fig. 5H; Supp. Fig. 6F).[45]

### Inflammatory monocytes are associated with improved prognosis across multiple tumor types

Our mouse data suggest that interferon-stimulated monocytes contribute to tumor control. To assess whether inflammatory monocytes are associated with improved prognosis in human tumors, we derived gene signatures of inflammatory monocytes and immunosuppressive macrophages (*Mmp12*+ macrophages) from our scRNAseq dataset to interrogate TCGA (Supplementary Table 1). We calculated the two signature scores for each patient, then subtracted the immunosuppressive macrophage score from the inflammatory monocyte score. By integrating both signatures into one score, we are comparing the relative tumor infiltration by these two myeloid populations across patient samples. This combined signature score was enriched in patients with improved outcomes in melanoma, bladder cancer, sarcoma, and many other malignancies (Fig. 6A, 6B). We next compared the performance of this signature with previously published prognostic signatures and the Hallmark IFNγ response signature.[46,47] Most prognostic signatures are highly correlated because they capture shared biology. Although the hazard ratios for death were similar for all prognostic signatures, our signature showed the lowest correlation with the others, suggesting it captures a potentially unique biology (Fig. 6C, 6D). To test if inflammatory monocytes are associated with improved response to immune checkpoint inhibitors, we combined published single-cell RNA-seq datasets from melanoma patients who received ICI.[48,49] Our inflammatory monocyte signature was significantly enriched in myeloid cells in pre-treatment samples from melanoma patients who responded to ICI therapy compared with non-responders (Fig. 6E, Supp. Fig. 7A, 7B). These data support the clinically relevant role of inflammatory monocytes for improved tumor control.

**Figure 6.**
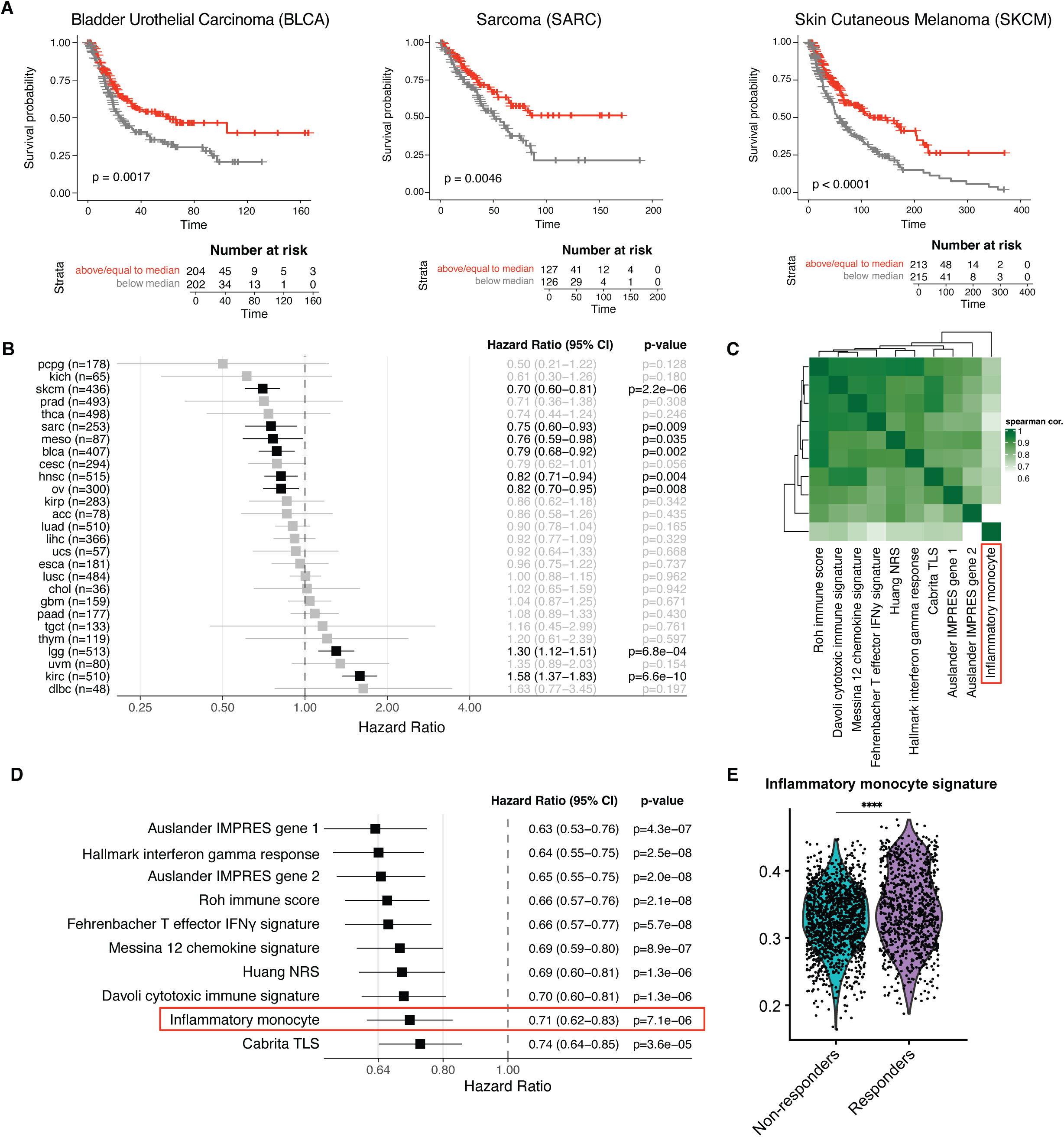
Inflammatory monocytes are associated with improved outcomes in cancer patients. **(A)** Survival analysis from TCGA dividing patients by median expression of inflammatory monocyte minus immunosuppressive macrophage signature score. Kaplan-Meier survival plots for bladder cancer (left), sarcoma (middle), and melanoma (right) shown. **(B)** Hazard ratio of death for all cancer subtypes from TCGA using our inflammatory monocyte minus immunosuppressive macrophage signature score, statistically significant results in black. **(C)** Spearman correlation coefficient matrix for indicated signatures across all patients in TCGA analysis. Inflammatory monocyte minus immuno-suppressive macrophage signature highlighted in red box. **(D)** Hazard ratio for death for all cancer subtypes from TCGA for different signatures. Inflammatory monocyte minus immunosuppressive macrophage signature highlighted in red box. **(E)** Inflammatory monocyte signature score for each myeloid cell isolated from pre-treatment biopsies of melanoma patients who received immune checkpoint inhibitor therapy, divided into responders and non-responders. Single-cell data combined from Sade-Feldman *et al.* and Pozniak *et al.* **** p < 0.0001

## DISCUSSION

Loss of antigen presentation is one mechanism by which tumor cells evade T cell-mediated killing, highlighting the need for new immunotherapy approaches to control these tumors. Using genetically modified mouse tumor cells, we demonstrated that CD40 agonist therapy can control *B2m*-null tumor growth, but unexpectedly this effect requires both CD8^+^ T cells and NK cells. Single-cell RNA-seq identified an increase in IFNγ production by these populations, which we confirmed *ex vivo*. IFNγ signaling in host cells, but not tumor cells, was required for the efficacy of CD40 agonist therapy. We further demonstrated that IFNγ acts on tumor-infiltrating monocytes to repolarize them toward a proinflammatory state. We derived an inflammatory monocyte signature that identifies patients with better prognosis across a variety of cancers and is enriched in melanoma patients who respond to immune checkpoint inhibitors. Together, these results suggest that monocyte-directed therapies may represent a promising strategy to control tumor growth in the setting of impaired antigen presentation.

We and others have established a critical role for IFNγ in the efficacy of CD40 agonist therapy. The loss of efficacy of CD40 agonist in *Ifngr1*^-/-^ mice suggests that the mechanism of action is dependent on IFNγ. Our data further argue against a model in which IFNγ acts directly on tumor cells to induce cell death.[41,44] Interestingly, IFNγ is also important for IL-2 mediated control of *B2m*-null B16 tumors, suggesting shared mechanisms of action.[5] IFNγ can be produced by many different immune effector cells, including CD4^+^, CD8^+^ T cells, and innate lymphocytes, such as NK cells. Our cell depletion and *ex vivo* cytokine studies suggest that production of IFNγ by both CD8^+^ T cells and NK cells is important, although we cannot rule out direct tumor cell killing by NK cells or CD8^+^ T cells.[5,12] However, the complete inhibition of CD40 agonist efficacy by IFNγ blockade argues against direct cytotoxicity. These data also help to explain how, despite the small number of neoantigen-specific CD8^+^ T cells found in most tumors, they are still critical for tumor control and correlate with patient outcomes. Our data support the hypothesis that IFNγ production by CD8^+^ T cells may be more important for tumor control than tumor antigen recognition and direct cytotoxicity.

Our depletion studies suggest that CD4^+^ T cells are dispensable in this model, despite evidence from other ICI-resistant and *B2m*-null models that they can be activated to control tumor growth.[5,11] Given the importance of IFNγ in our model and the observation that CD40 agonist increases CD4^+^ T cell infiltration of tumors and IFNγ production by CD4^+^ T cells (Fig. 3D, 4D), the lack of impact of CD4^+^ T cell depletion was particularly striking. We hypothesize that despite their *ex vivo* production of IFNγ, effector CD4^+^ T cells are either not major contributors to IFNγ in the TME or not spatially localized near infiltrating monocytes to stimulate them in our model.

Although NK cells, CD4^+^ T cells, CD8^+^ T cells, and yδ T cells have been shown to control *B2m*-null tumors,[5,10,12,13] our data support a previously underappreciated role for monocytes in *B2m*-null tumor control. The ability of many different cell types to control *B2m*-null tumors suggests that there may be multiple strategies to overcome resistance to ICI. The clinical challenge will be to determine which patients require which immune cells to achieve effective tumor control and to identify cell-targeting strategies that avoid the systemic toxicity of CD40 agonists. Supporting a role for inflammatory monocytes, CIBERSORTx analysis of pretreatment human melanoma biopsies found that in patients with loss of heterozygosity in *B2M*, ICI responders had higher monocyte numbers in tumors than non-responders.[5] In addition, inflammatory monocytes were enriched in anti-PD-1 responsive syngeneic tumor models compared with non-responsive models.[50] The precise mechanisms by which inflammatory monocytes control *B2m*-null tumor growth remain to be defined.

In conclusion, we identified a role for CD8^+^ T cells and NK cells in controlling *B2m*-null tumor growth through production of IFNγ, which promotes inflammatory monocyte-mediated tumor control. Our findings suggest that therapeutic strategies aimed at activating inflammatory monocytes may improve outcomes for patients who do not respond to T cell-directed cancer immunotherapies.

## METHODS

### Mice and cell lines

Female wild-type C57BL/6J (strain # 000664), *Tcra^-/-^*(B6.129S2-*Tcra^tm1Mom^*/J; strain # 002116) and *Ifngr1*^-/-^ (B6.129S7-*Ifngr1^tm1Agt^*/J; strain # 003288) mice were purchased from The Jackson Laboratory. NSG (NOD.Cg-*Prkdc^scid^Il2rg^tm1Wjl^*/SzJ; Jackson strain # 005557) mice were obtained from the UNC Chapel Hill Preclinical Research Unit. Parental and GVAX (GM-CSF expressing) B16F10 melanoma cell lines were provided by G. Dranoff. CRISPR/Cas9-mediated mutation of the *B2m*, *H2-K1*, *H2-D1*, and *Ifngr1* loci in B16.F10 and/or MC38 cell lines was accomplished by transient transfection with a plasmid expressing Cas9 (PX459, Addgene) and guide RNAs targeting *B2m* (CACCGACAAGCACCAGAAAGACCA), *H2-K1* (CACCGTAGCCGACTTCCATGTACCG), *H2-D1* (CACCGTAGCCGACAGAGATGTACCG), or *Ifngr1* (CACCCGACTTCAGGGTGAAATACG). All transfected cells were puromycin selected and verified by flow cytometry after co-culturing with 100ng/mL IFNγ for 48 hours to upregulate MHC-I or PD-L1 (Supp. Fig. 1A, 2A, 5B). All animal procedures were compliant with ethical guidelines approved by the University of North Carolina-Chapel Hill Institutional Animal Care and Use Committee.

### *In vivo* tumor studies

For all tumor studies, mice were subcutaneously implanted with 1*10^6^ tumor cells. Where indicated, mice were vaccinated with an equivalent number of irradiated (3,500 cGy) GVAX cells on days 1 and 4 post-implantation of the primary tumor. Tumors were measured every 2-3 days. Treatment with 100µg CD40 agonist (clone FGK4.5, BioXCell) or 100µg of isotype-matched control antibody (clone 2A3, BioXCell) was administered intraperitoneally every 2-3 days beginning once the average tumor volume across all experimental groups eclipsed 100mm^3^. Where indicated, tumor-bearing mice were treated with 100µg ɑPD-1 (clone 29F.1A12, BioXCell) or 100µg of isotype-matched control antibody (clone 2A3, BioXCell) on days 6, 9, and 12 post-implantation. For antibody-mediated depletion studies, mice were intraperitoneally injected with 200µg ɑCD19 (clone 1D3, BioXCell), 200µg ɑTCR𝛄δ (clone UC7-13D5, BioXCell), 200µg ɑCD8a (clone 2.43, BioXCell), ɑCD4 (clone GK1.5, BioXCell), and/or ɑNK1.1 (clone PK136, BioXCell) or 200µg of appropriate isotype-matched control antibodies (clone 2A3, clone HRPN, clone LTF-2, clone C1.18.4, BioXCell) the day prior to tumor implantation and every 2-3 days post-implantation. For cytokine neutralization experiments, tumor-bearing mice were treated with 500µg ɑIFNγ (clone XMG1.2, BioXCell) or 500µg isotype-matched control antibody (clone HRPN, BioXCell) via intraperitoneal injection on days 6, 9, and 12 post-tumor implantation.

### Tumor dissection and cell isolation

Tumor-bearing mice were euthanized via CO_2_ and cervical dislocation, tumors were excised and placed in digestion media (RPMI-1640 (Gibco) + 2% FBS (Gemini Bio-Products) + 2mg/mL collagenase P (Sigma) + 50mg/mL DNase I (Sigma)) and minced using dissection scissors. Minced tumors were digested at 37°C for 30 minutes. Fully-digested tumors were vortexed, filtered into a fresh conical through a 70µm cell-strainer, and washed 1-2x in cold R2 (RPMI-1640 (Gibco) + 2% FBS (Gemini Bio-Products)). Single-cell suspensions of digested tumors were then purified for immune cell populations using MACS (CD45+ or CD11b+) bead-based positive selection (Miltenyi) for downstream applications.

### *Ex vivo* cytokine production

For monocyte *ex vivo* cytokine production assays, CD11b^+^ cells were purified from tumors using CD11b+ MACS and 500,000 CD11b^+^ cells were plated in non-TC treated 24-well flat-bottom plates. All wells received 1X Protein Transport Inhibitor Cocktail (eBioscience) containing Brefeldin A and Monensin to sequester cytokines intracellularly. Stimulated monocyte wells were supplemented with 100ng/mL LPS (Sigma). After 18 hours of *ex vivo* culture, adherent cells were harvested by vigorously washing the surface of each well with cold PBS (Gibco) + 2mM EDTA (Gibco). Monocytes were analyzed for intracellular cytokines by flow cytometry.

For tumor-infiltrating lymphocyte (TIL) *ex vivo* experiments, CD45^+^ cells were purified from tumors, counted, and 0.5-1*10^6^ cells plated in TC-treated 24-well flat-bottom plates. Unstimulated wells were supplemented with 1X Protein Transport Inhibitor Cocktail (eBioscience) while stimulated wells received 1X Cell Stimulation Cocktail (plus protein transport inhibitors) (eBioscience). After 6 hours of *ex vivo* culture, intracellular IFNγ was detected by flow cytometry.

### Murine single-cell RNA-seq library preparation and analysis

*B2m*-null B16F10 tumors were resected from C57BL6/J mice on day 14 after treatment with isotype control or CD40 agonist antibody on days 6, 9, and 12, as described above. TILs were isolated using CD45^+^ MACS beads per manufacturer instructions. Cells were counted and 250,000 TILs per mouse resuspended in blocking buffer (0.25ug TruStain FcX PLUS (Biolegend), 5% normal rat serum, 5% normal mouse serum) for 10 minutes. 0.5ug of TotalSeq-B hashtag anti-mouse antibodies (Biolegend, one hashtag per mouse in each treatment group) was added to the cells for 30 minutes. After washing twice, cells were pooled together from each experimental group, recounted, and loaded onto the Chromium Controller (10X Genomics) for a target recovery of 10,000-14,000 cells/lane. Gene expression and feature barcode libraries were generated per manufacturer’s instructions and sequenced at a 9:1 ratio on an Illumina NextSeq500 using a 75-bp kit with paired-end reads.

Sample demultiplexing, barcode processing, alignment, filtering, and UMI counting were performed using the Cell Ranger analysis pipeline (v.3.0.0). RNA and hashtag-oligo counts were subsequently used for downstream analysis. Data pre-processing, normalization, integration, and clustering was performed using the Seurat package (v.4.2.0) in R (v.4.2.1 “Funny-Looking Kid”). For each sample, cells with poor quality metrics were removed if they had fewer than 200 unique genes or greater than 10% mitochondrial counts. The resulting expression matrix contained 99,337 cells by 32,288 genes. Samples were demultiplexed using hashtag-oligos. Doublet cells were identified using hashtags and transcriptional profiles and then removed from downstream analyses. Signature scoring was performed using VISION (v.3.0.0), utilizing signatures from the Molecular Signature Database (MSigDB). Signatures were accessed through the msgidbr package (v7.5.1). Pseudobulk samples were created by aggregating the counts for each gene across all cells in each treatment group and genotype using functions from the custom package Rsc (v.0.0.900), after removing cells which were outliers by median absolute deviation (MAD) and considering only genes with greater than 10 total counts for each pseudobulk, using functions from scater (v.1.18.6) and Matrix.utils (v.0.9.8). DESeq2 (v.1.30.1) was used to normalize the pseudobulk counts matrix and perform differential gene expression. Pre-ranked gene set enrichment analysis was performed subsequently, using the Lightweight Iterative Geneset Enrichment in R (LIGER) tool (v2.0.1).

### Human single-cell RNA-seq analysis

Pre-processed human single-cell RNA-seq datasets of the tumor microenvironment were downloaded and split based on sample IDs (https://doi.org/10.48804/GSAXBN, GSE120575).[48,49] The sample-level single cell Seurat objects were normalized using SCTransform and integrated using the RPCA method. Next, we integrated the Seurat objects from both cohorts using the RPCA method. The integrated object was scaled and clustered using the default Louvain algorithm. Myeloid cells were subsetted based on canonical genes and reclustered to generate myeloid specific cell clustering. We calculated a per cell score for our inflammatory monocyte signature using UCell and tested for a difference in average signature score from pre-treatment samples between responders and non-responders using the Wilcoxon rank-sum test. Next, we performed differential gene expression analysis using the FindMarkers function in Seurat between all myeloid cells in pre-treatment samples from responders versus non-responders. We then performed GSEA to evaluate signature enrichment of our inflammatory monocyte signature.

### Human survival analysis

The Cancer Genome Atlas (TCGA) gene expression data and clinical metadata were downloaded from the cBioPortal website, selecting patient samples associated with “TCGA PanCancer Atlas Studies.” Signatures were scored using single sample GSEA, implemented in the GSVA package (v1.44.5), then z-scored using the scale function in R.

For the inflammatory monocyte versus immunosuppressive macrophage signature, we extracted the top 200 differentially expressed genes for the inflammatory monocyte or *Mmp12*+ macrophage clusters relative to all other cells in the single cell dataset, calculated a signature score for each patient sample, then calculated the z-score for each signature across samples. Because we wanted to capture the effect of inflammatory monocyte relative to immunosuppressive macrophages, we then calculated the difference between the inflammatory monocyte signature and the immunosuppressive macrophage signature. The final score was again z-scored across samples. Survival analysis was performed in R using the survival package (v3.5-0) and survival graphs were generated using the survminer package (v0.4.9) in R. P-values for statistical tests comparing discrete variables are from the Log-rank test; p-values for statistical tests comparing continuous variables are from the Cox Proportional Hazard test.

### Statistical analysis

Where possible, mice in tumor studies were randomized before beginning treatment. For all tumor growth and *ex vivo* experiments, 5-10 individual mice per experimental group were used to guarantee sufficient statistical power. Experimental results were analyzed via unpaired two-sided Student’s *t*-tests using GraphPad Prism 10. Outlier data points in mouse experiments were identified and removed using Rout outlier tests executed with default Prism parameters. Statistical significance was signified with asterisks that correspond to the following p values: ns, not significant (p > 0.05), *p ≤ 0.05, **p ≤ 0.01, ***p ≤ 0.001 and ****p ≤ 0.0001.

## Data and code availability

Data are available in a public, open access repository. All code used to process the single cell RNA-seq data can be found on GitHub at https://github.com/amyh25/b2m-cd40-quarto. Raw and processed single cell RNA-seq data have been deposited in the Gene Expression Omnibus under accession number GSE240148 and to the Broad Institute Single Cell Portal under accession number SCP2033. Gene sets analyzed with GSEA are available through the Broad Institute Molecular Signatures Database or have been included in the GitHub.

See Supplemental Methods for additional details.

## Authors Contributions

Conception and design: WNH, AHS, BCM

Acquisition of data: MPZ, EKC, RA, WLC, AGB, KJV, YC, HW, MF, SAW, JRL, TV, SMP, BCM

Analysis and interpretation of data: MPZ, AYH, EKC, RA, WLC, AGB, YC, VZ, EJR, TV, SP, WH, DRS, BCM

Writing, review, and/or revision of the manuscript: MPZ, AYH, EKC, RA, KJV, SAW, TV, WNH, AHS, BCM

Administrative, technical, or material support: JC, SAW, MF

Study supervision: PKS, WNH, AHS, BCM

## Acknowledgements

The UNC Flow Cytometry Core Facility (RRID:SCR_019170) is supported in part by P30 CA016086 Cancer Center Core Support Grant to the UNC Lineberger Comprehensive Cancer Center. Colony management services were performed by the Colony Management Core within the Division of Comparative Medicine as well as the UNC Lineberger Preclinical Research Unit at the University of North Carolina at Chapel Hill which is supported in part by an NCI Center Core Support Grant (CA16086) to the UNC Lineberger Comprehensive Cancer Center. This work was presented at the 2023 AACR National Meeting.[1]

## List of Abbreviations

B2m: beta-2 microglobulin
MHC-I: major histocompatibility complex I
TNF: tumor necrosis factor
TAM: tumor-associated macrophage
IFN: interferon
LPS: lipopolysaccharide
agCD40: CD40 agonist
scRNAseq: single-cell RNA-sequencing
TME: tumor microenvironment
TIL: tumor-infiltrating lymphocyte

---

**Supplementary Figure 1.**
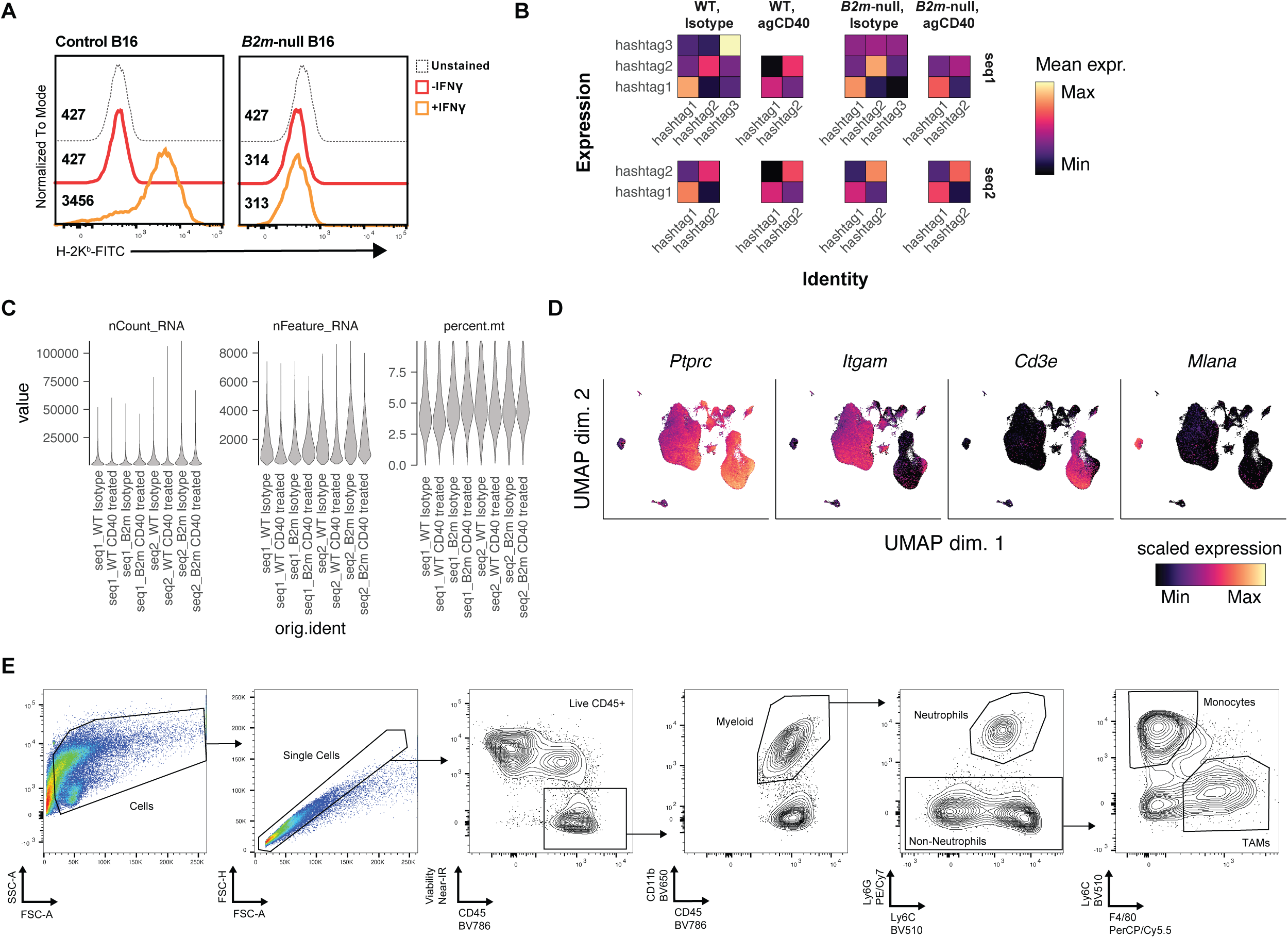
**(A)** Flow cytometry analysis of H-2K^b^ expression on control or *B2m*-null B16 tumor cells with or without 48 hours IFNγ stimulation. Geometric mean fluorescence intensity shown. Full-minus one (FMO) staining control shown. **(B)** Heatmaps demonstrating average expression of the indicated hashtag sequence within cells assigned to each identity. **(C)** Quality control metrics for the different samples sequenced for single-cell RNA-seq. **(D)** Expression of the indicated genes on the UMAP across all samples and cell types. **(E)** Flow cytometry gating strategy for intratumoral myeloid populations.

**Supplementary Figure 2.**
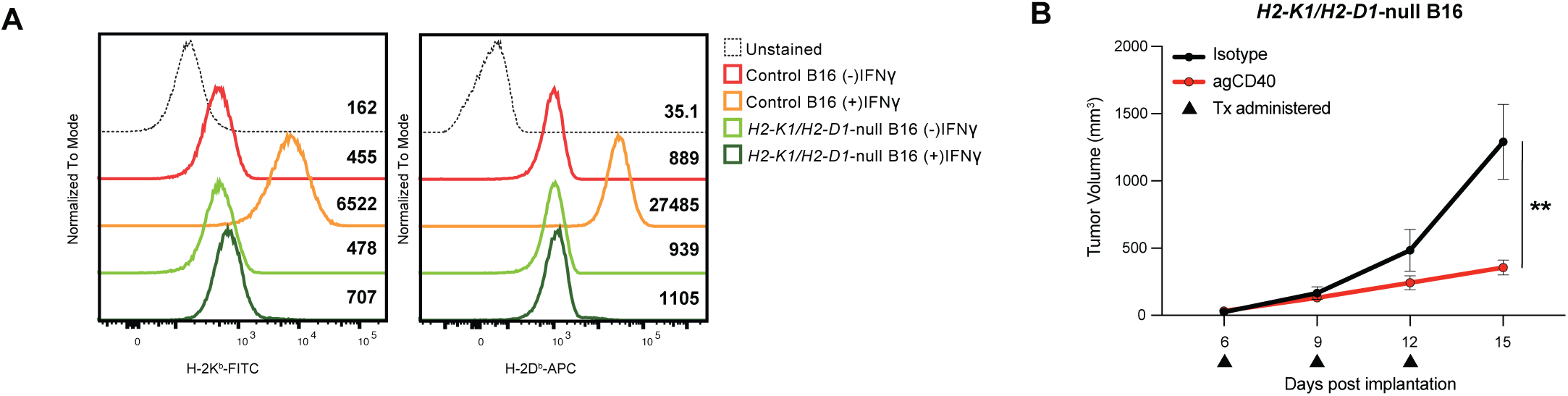
**(A)** Flow cytometry analysis of H-2K^b^ and H-2D^b^ expression on control or *H2-K1/H2-D1*-null B16 tumor cells with or without 48 hours IFNγ stimulation. Geometric mean fluorescence intensity shown. Full-minus one (FMO) staining control shown. **(B)** Tumor growth curve of *H2-K1/H2-D1*-null B16 tumor cells implanted into C57BL/6J mice and treated with isotype or agCD40 antibodies on days 6, 9, and 12 post-implantation. ** p < 0.01.

**Supplementary Figure 3.**
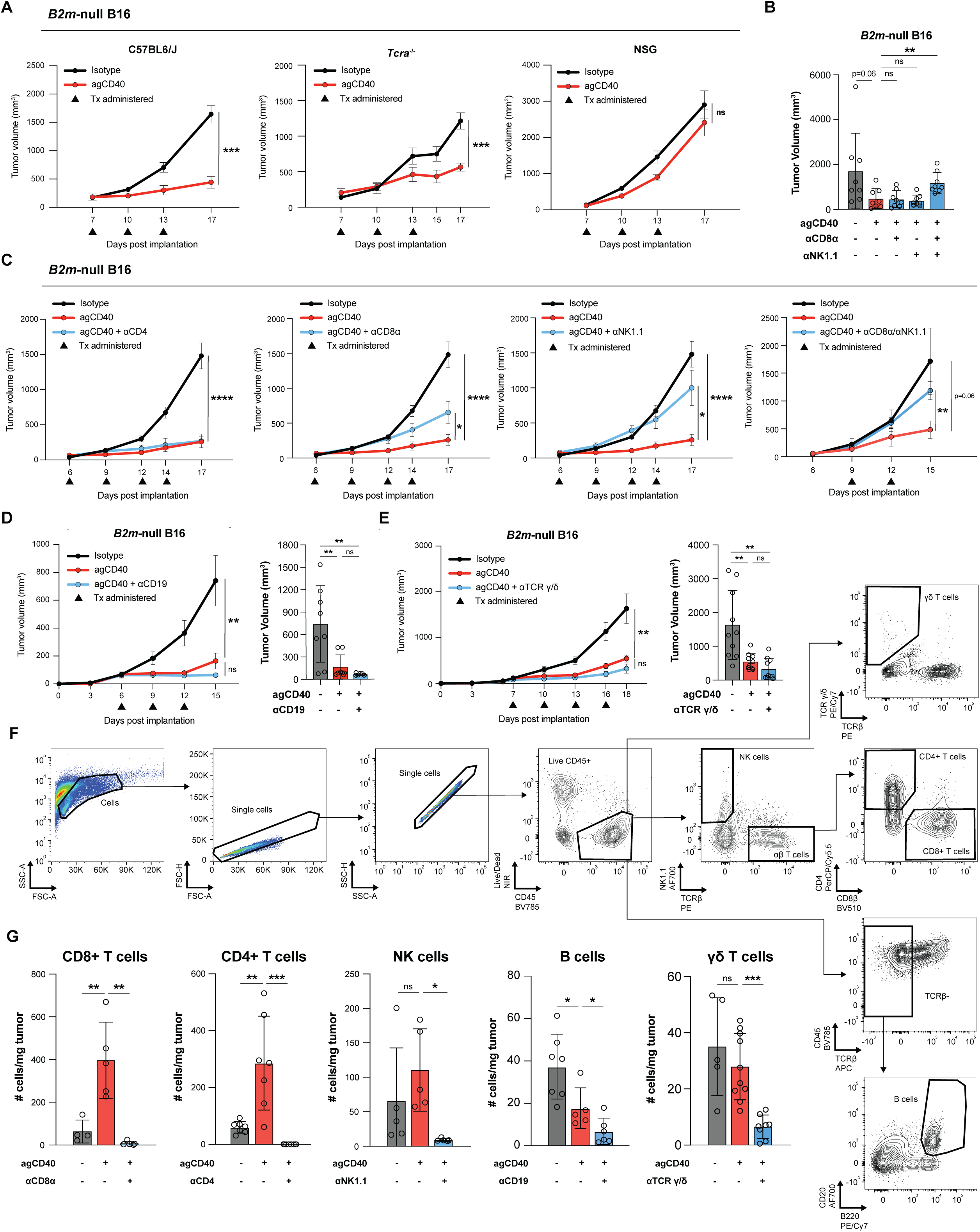
**(A)** Growth curves of *B2m*-null B16 tumors implanted into mice of the indicated genotypes and treated with either CD40 agonist or isotype control antibodies. **(B-E)** Growth curves and final tumor volumes of *B2m*-null B16 tumors implanted into C57BL6/J wild-type mice depleted of the indicated cell types and then treated with CD40 agonist or isotype control antibodies. **(F)** Flow cytometry gating strategy for intratumoral lymphocyte quantification. **(G)** Quantification of the number of immune cells per milligram tumor in the indicated depletion experiments. * p < 0.05; ** p < 0.01; *** p < 0.001; **** p < 0.0001; n.s. not significant.

**Supplementary Figure 4.**
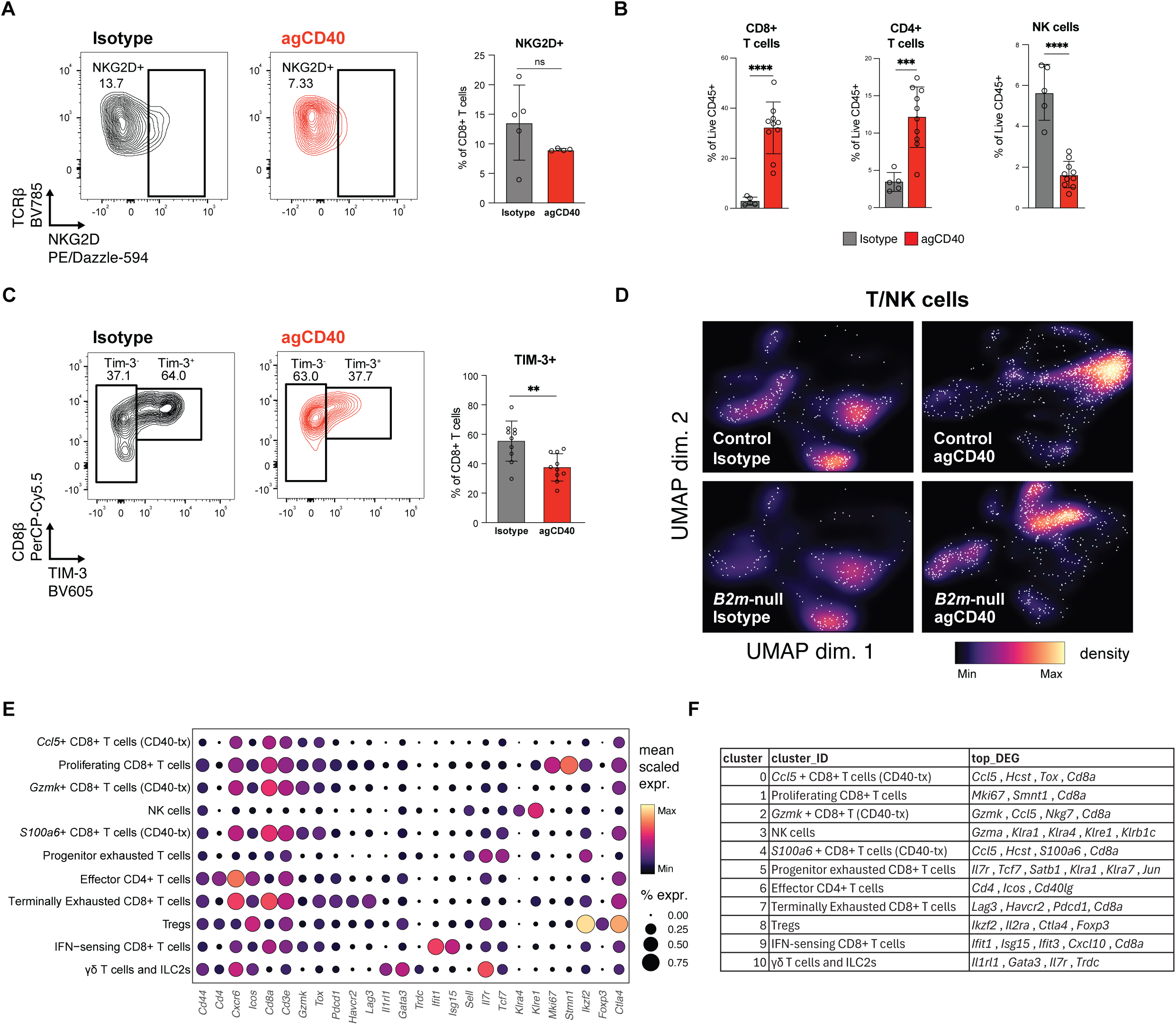
**(A)** NKG2D expression on CD8^+^ T cells from *B2m-null* B16 tumors treated with isotype control (n = 5) or agCD40 (n = 4) antibody. Representative flow plots (left) and quantification (right). **(B)** Flow cytometry quantification of CD8^+^ T cell, CD4^+^ T cell, and NK cell populations as a percentage of Live CD45^+^ cells in *B2m-null* B16 tumors treated with isotype control or agCD40 antibody. Representative results from at least 3 independent experiments. **(C)** TIM-3 expression on CD8^+^ T cells from *B2m-null* B16 tumors treated with isotype control or agCD40 antibody. Representative flow plots (left) and quantification (right) from one of two independent experiments. **(D)** Galaxy plot displaying cell density of T/NK cells subclustered from scRNAseq analysis of control and *B2m-null* B16 tumors treated with agCD40 or isotype-control antibody. **(E)** Dot plot showing indicated mean gene expression (color) and percentage of cells expressing the gene (size) for each of the T/NK cell subclusters. **(F)** Table of T/NK cell subclusters and top differentially expressed genes within each cluster. ** p < 0.01; *** p < 0.001; **** p < 0.0001; n.s. not significant.

**Supplementary Figure 5.**
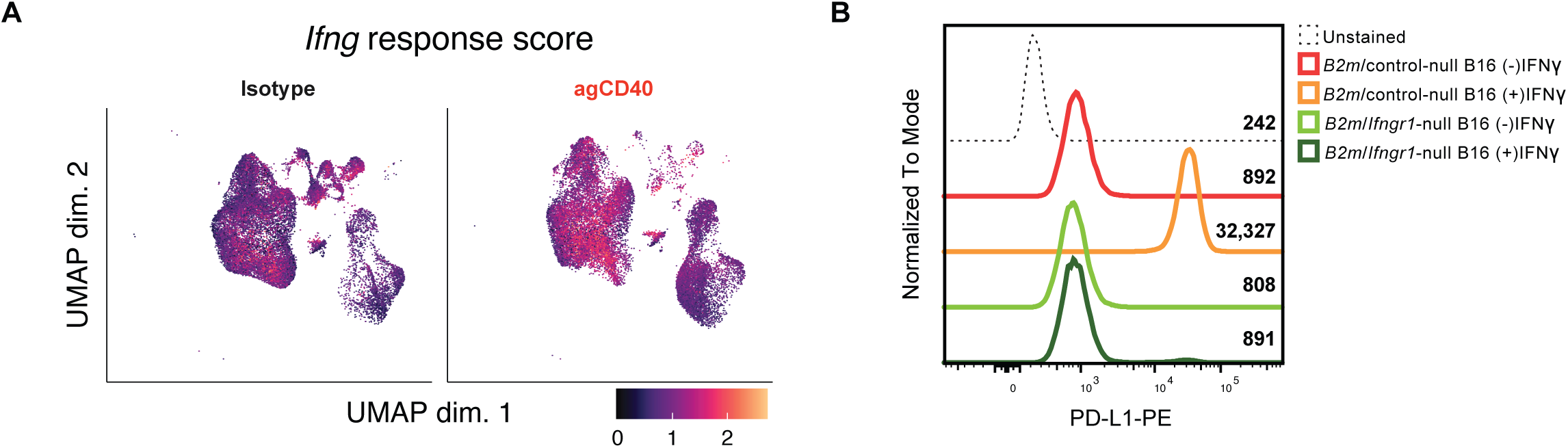
**(A)** Expression of the Hallmark IFNγ response signature on the UMAP showing myeloid cells and T/NK cell clusters. **(B)** Flow cytometry analysis of PD-L1 expression on *B2m-null/control* or *B2m-null/Ifngr1-null* B16 tumor cells with or without 48 hours IFNγ stimulation. Geometric mean fluorescence intensity shown. Full-minus one (FMO) staining control also shown.

**Supplementary Figure 6.**
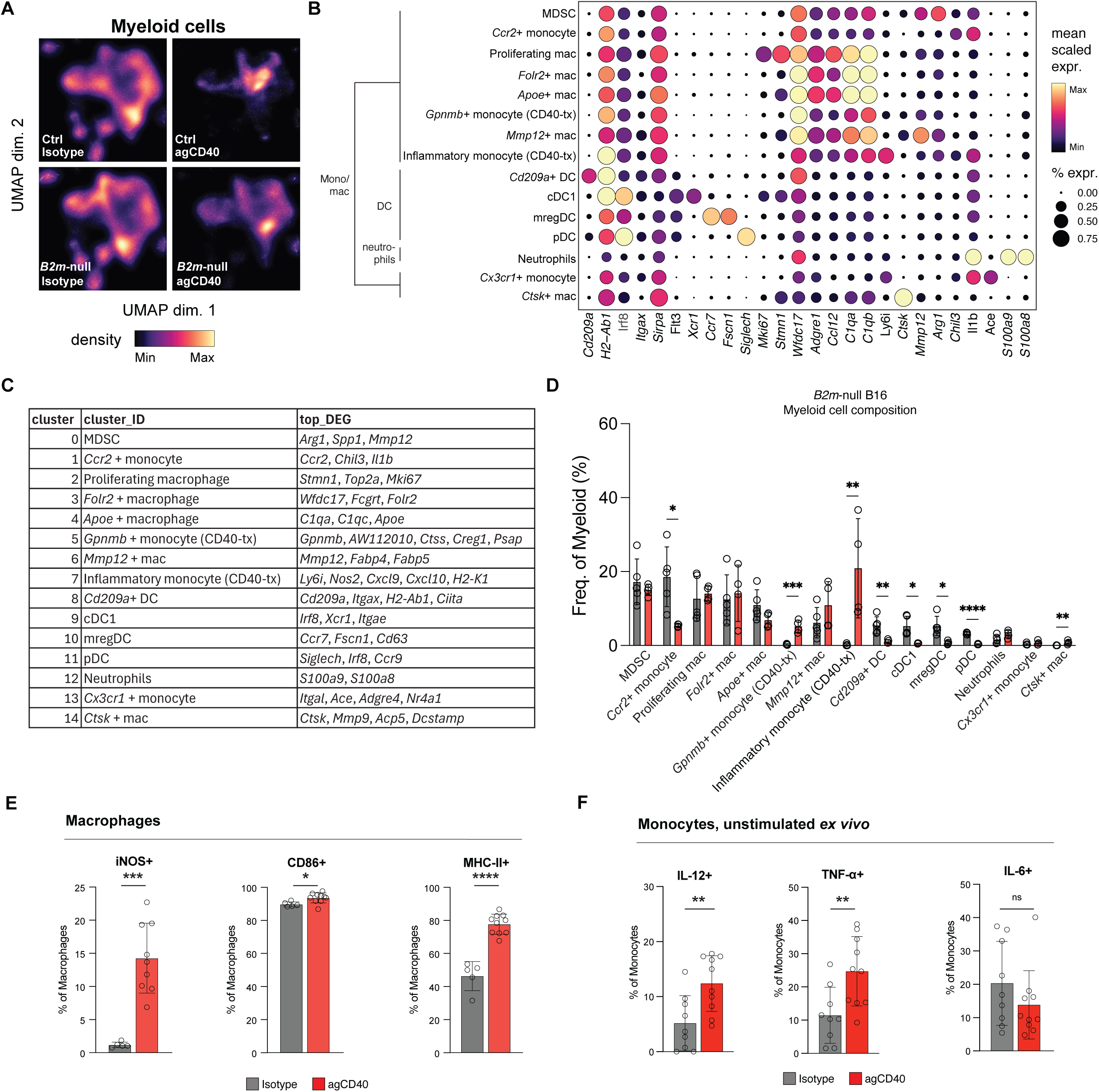
**(A)** Density plot in myeloid cluster UMAP space for each treatment condition. **(B)** Dot plot showing indicated mean gene expression (color) and percentage of cells expressing the gene (size) for each of the myeloid cell subclusters. **(C)** Table of myeloid cell subclusters and top differentially expressed genes within each cluster. **(D)** Quantification of each cell cluster from scRNAseq analysis as a frequency of total immune cells. **(E)** Frequency of iNOS+, CD86+, and MHC-II+ tumor-associated macrophages from isotype control- or agCD40-treated *B2m*-null B16 tumors. **(F)** Frequency of IL-12p40+, TNFα+, and IL-6+ monocytes isolated from isotype control- or agCD40-treated *B2m*-null B16 tumors and plated without stimulation *ex vivo* for 18h. * p <0.05; ** p < 0.01; *** p < 0.001; **** p < 0.0001; ns not significant.

**Supplementary Figure 7.**
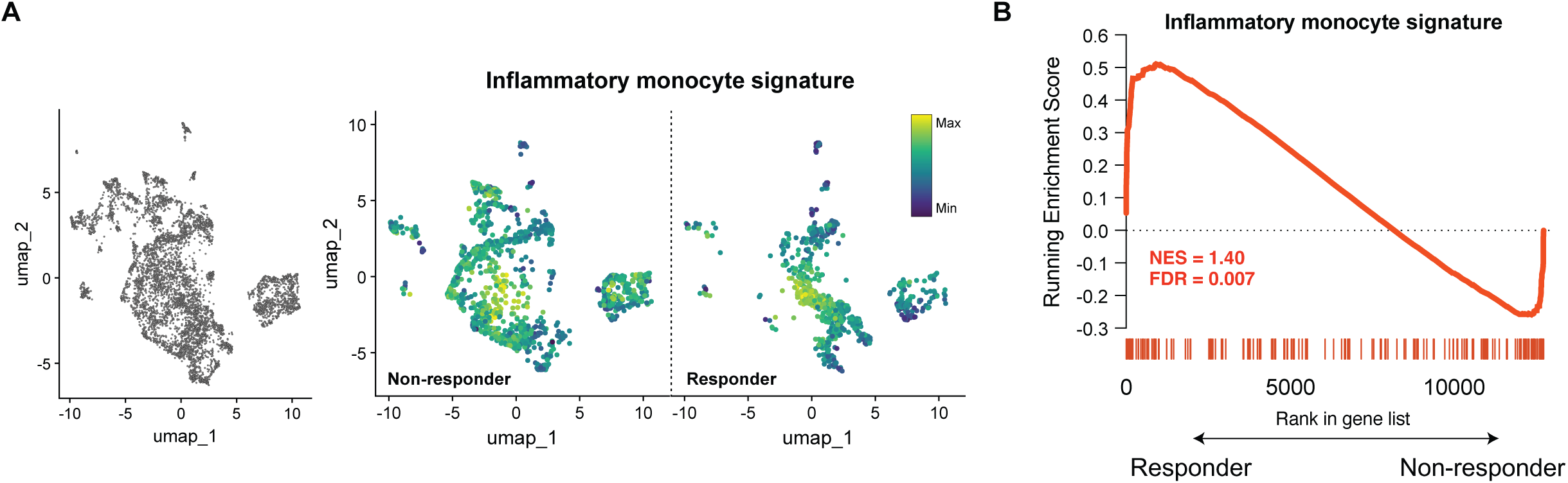
**(A)** UMAP and Inflammatory monocyte signature score in individual myeloid cells (non-responder left, responder right) from human myeloid cell analysis. **(B)** GSEA of the inflammatory monocyte signature in the ranked list of genes differentially expressed in pre-treatment myeloid cells between responders and non-responders. * p <0.05; ** p < 0.01; *** p < 0.001; **** p < 0.0001.

