## Supplemental Table 1 for "Harnessing Inflammatory Monocytes to Overcome Resistance to Anti-PD-1 Immunotherapy"

**Supplementary Table 1**

| Inflammatory_monocytes | Immunosuppressive_macrophages |
|---|---|
| Ly6i | Mmp12 |
| AA467197 | Fabp5 |
| Prdx5 | Fabp4 |
| Cfb | Gpnmb |
| Sod2 | Ctsl |
| Lyz2 | Ftl1 |
| Clec4e | Lgals3 |
| Nos2 | Fth1 |
| C3 | Atp6v0d2 |
| Clec4n | Gdf15 |
| Saa3 | Prdx1 |
| H2-Ab1 | Hmox1 |
| H2-Eb1 | Mmp13 |
| Cd74 | Esd |
| Upp1 | Cd63 |
| Cxcl9 | Spp1 |
| Tgfbi | Gpr137b |
| Fpr2 | Cd68 |
| Fbxl5 | Ndrg1 |
| H2-Aa | Lpl |
| Pla2g7 | Ctsb |
| Ctsc | Cd36 |
| Ly6a | Pmp22 |
| Ass1 | Dnmt3a |
| Slc7a11 | Il1rn |
| Acod1 | Trem2 |
| Cxcl10 | Atp6v1a |
| Cybb | Cstb |
| Ifi30 | Hilpda |
| Lyz1 | Pgam1 |
| Gbp2 | Aldoa |
| Mcemp1 | Ctsd |
| Il1b | Creg1 |
| Ifitm3 | Basp1 |
| Cd14 | Anxa1 |
| Bst1 | Bnip3 |
| Ptgs2 | Mt1 |
| Aif1 | Lipa |
| Plac8 | Plin2 |
| H2-DMb1 | Gde1 |
| Clec12a | Igf1 |

Blvrb
Marcksl1
Tspo
Slamf8
Pilra
Cxcl2
Iigp1
Klra2
H2-DMa
Pid1
Atox1
Clec5a
Ccrl2
Slc7a2
Cxcl16
Prdx6
Nampt
Pirb
Capg
Cstb
Calhm6
Gm15056
Glrx
Slc16a3
Grina
Txn1
Magohb
Cebpb
Tgm2
Ms4a6c
Cyba
Zbp1
F10
S100a11
Gda
Clec4d
Bcl2a1a
Slpi
Lpcat2
Acsl1
Psme2
Lst1
Dram1
Mif
Osbpl8
Samd8
Bhlhe41
Timp2
Serpinb6a
Psmb6
Plk2
Adam8
Slc2a1
Clec4d
Psmd8
Akr1a1
Atp6v0c
Tmem65
Ccl8
Vat1
Tpi1
Ero1l
Plau
Card19
Ldha
Ninj1
Abca1
Atp6v1c1
Ctsz
Gapdh
Rnh1
Pkm
Cd9
Lhfpl2
Gng11
Gclm
Ctsk
Eif4ebp1
C1qc
Fam129b
Il7r
Ctsa
Vim
C1qb
Rilpl2
Tmem189

Ms4a4c
Prdx1
Pnp
Tlr2
Ehd1
Fcer1g
Psmb10
Fth1
Hck
Msr1
Itgb2
Ms4a6d
Pkm
Ctsz
Bcl2a1b
Gpr141
Naaa
Ier3
Rab32
Inhba
Cfp
Fn1
Msrb1
Cd40
Samhd1
Arg1
Mmp14
Vim
Plaur
Esd
Tnfaip2
Fgr
Cnih4
Smpdl3b
Tyrobp
Atp6v0c
Tapbp
Csf1r
Gpx1
Procr
Pgd
Fcgr4
Cdkn1a
Bsg
Sgk1
Capg
Gyg
Sat1
Blvrb
Pgk1
Psap
Atf3
Por
Pf4
Lrp12
Mt2
Nceh1
Hk2
Gstm1
Lgmn
Nos2
Pld3
Gabarap
Fnip2
Rassf8
Fblim1
Cpeb4
Bnip3l
Mtss1
Vcam1
Slc30a1
Rnf128
Anxa4
Atp6v1g1
Sqstm1
Sdcbp
Ncoa4
Srxn1
Grn
Litaf
Cyba
Lamp1
Apoe
Ftl1-ps1
Lgals1
Lat2

Psmb8
Itgam
Spi1
Tpi1
Sdc4
Rnf19b
Isg15
Alas1
Alox5ap
Ncf4
Mcub
Tma16
Emp3
Mif
Cd52
Ly6c2
App
Socs3
Nrp2
Ifitm2
Gapdh
Dusp1
Lair1
Gsn
Nlrp3
Psma7
Creb5
Fcgr1
Plbd1
Sh3bgrl3
Snx10
Ninj1
Aldoa
Ptpn6
Gbp4
Nupr1
Stat1
Pomp
Sdc3
Srgn
Ctss
Ggh
Cd274

Slc6a8
Ccl12
Soat1
Tent5c
Hexa
Mgst1
Atp6v0e
Emp1
Pdpn
Gadd45a
Bri3
Sh3glb1
Slc7a2
Cyb5a
Vdac1
Gla
Vegfa
Slamf7
Anpep
Vdac2
Vapa
Slc48a1
Cxcl16
C1qa
Hist1h2bc
Gna13
Rgl1
Syngr1
Ankrd37
Dstn
Atp6v0b
Gng2
Arhgap10
Grb2
Ap3s1
Myo5a
Tpp1
Abcg1
Dhrs3
Npc2
Gpi1
Rgs1
Ccl9

| | |
|---|---|
| Irf7 | 0610012G03Rik |
| Slc11a1 | Rcbtb2 |
| Sirpb1c | Mxi1 |
| Mvp | Arl8b |
| Slc7a8 | Anxa5 |
| Ly6e | Rsad2 |
| Fcgr2b | Sdc1 |
| Tppp3 | Atp6v1b2 |
| Unc93b1 | Slc27a1 |
| N4bp1 | Rhoc |
| Hk3 | Mpc1 |
| Ptpn1 | Ctss |
| Fos | Hebp1 |
| Prkcd | Tmem106a |
| S100a4 | Ms4a7 |
| Ncf1 | Ugp2 |
| Vamp8 | Vcl |
| Card19 | Plekho1 |
| Ifi27l2a | Rnasek |
| Rrbp1 | Aph1c |
| Ccr1 | Plxna1 |
| Eno1 | Abcc5 |
| Ltb4r1 | Rragc |
| Cyfip1 | Hspa1a |
| Pgam1 | Aprt |
| Ppt2 | Ccl6 |
| BC028528 | Rala |
| Tmbim4 | Flrt2 |
| Scimp | C3ar1 |
| Gm4951 | Pdgfa |
