## Supplemental Table 2 for "Harnessing Inflammatory Monocytes to Overcome Resistance to Anti-PD-1 Immunotherapy"

| Supplementary Table 2 Antibodies | | | | |
|---|---|---|---|---|
| **Target** | **Clone** | **Company** | **Fluorophore** | **Dilution** |
| Arg1 | A1exF5 | eBioscience | AF700 | 1:200 |
| CD11b | M1/70 | BD Biosciences | BUV395 | 1:800 |
| CD11b | M1/70 | BioLegend | BV421 | 1:400 |
| CD11b | M1/70 | BioLegend | BV650 | 1:400 |
| CD19 | 1D3 | BD Biosciences | PE | 1:400 |
| CD206 | C068C2 | BioLegend | AF594 | 1:50 |
| CD206 | C068C2 | BioLegend | FITC | 1:200 |
| CD206 | C068C2 | BioLegend | BV421 | 1:200 |
| CD3ε | 145-2C11 | BioLegend | BV421 | 1:100, 1:400 |
| CD4 | GK1.5 | BioLegend | PerCP/Cy5.5 | 1:200, 1:400 |
| CD4 | GK1.5 | BioLegend | FITC | 1:400 |
| CD44 | IM7 | BioLegend | APC/Cy7 | 1:200 |
| CD45 | 30-F11 | BioLegend | BV785 | 1:400 |
| CD45 | 30-F11 | BioLegend | PerCP/Cy5.5 | 1:400 |
| CD45.2 | 104 | BioLegend | BV605 | 1:400 |
| CD86 | GL-1 | BioLegend | PE/Cy7 | 1:400 |
| CD86 | GL-1 | BioLegend | BV785 | 1:400 |
| CD8α | 53-6.7 | BioLegend | AF594 | 1:200 |
| CD8β | YTS156.7.7 | BioLegend | BV510 | 1:200, 1:400 |
| CD8β | YTS156.7.7 | BioLegend | PerCP/Cy5.5 | 1:200 |
| F4/80 | BM8 | BioLegend | AF594 | 1:400 |
| F4/80 | BM8 | BioLegend | APC/Cy7 | 1:200 |
| F4/80 | BM8 | BioLegend | PerCP/Cy5.5 | 1:200 |
| F4/80 | T45-2342 | BD Biosciences | BUV563 | 1:400 |
| H-2Db | KH95 | BioLegend | APC | 1:100 |
| H-2Kb | AF6-88.5 | BioLegend | FITC | 1:100 |
| IFNγ | XMG1.2 | BioLegend | PE | 1:100, 1:400 |
| IL-12/23-p40 | C15.6 | BioLegend | PE/Dazzle 594 | 1:100 |
| IL-6 | MP5-20F3 | eBioscience | FITC | 1:100 |
| iNOS | W16030C | BioLegend | PE | 1:400 |
| Ly6G | 1A8 | BioLegend | PE/Cy7 | 1:400 |
| Ly6C | HK1.4 | BioLegend | BV510 | 1:400 |
| Ly6C | HK1.4 | BioLegend | BV605 | 1:400 |
| Ly6C | HK1.4 | BioLegend | FITC | 1:400 |
| MHCII | M5/114.15.2 | BioLegend | PerCP/Cy5.5 | 1:400 |
| MHCII | M5/114.15.2 | BioLegend | PE | 1:400 |
| NK1.1 | PK136 | BioLegend | AF700 | 1:400 |
| NK1.1 | PK136 | BioLegend | PE | 1:400 |
| NKG2D | CX5 | BioLegend | PE/Dazzle 594 | 1:200 |
| NKp46 | 29A1.4 | BioLegend | PE/Cy7 | 1:100, 1:200 |
| PD-1 | 29F.1A12 | Biolegend | BV421 | 1:200 |

| | | | | |
|---|---|---|---|---|
| PD-1 | APC | Biolegend | APC | 1:200 |
| TCRβ | H57-597 | BioLegend | PE | 1:400 |
| TCRβ | H57-597 | BioLegend | BV785 | 1:400 |
| TCRβ | H57-597 | BioLegend | APC | 1:400 |
| TCR$\gamma\delta$ | GL3 | BioLegend | PE/Cy7 | 1:400 |
| Tim-3 | RMT3-23 | BioLegend | BV605 | 1:100 |
| TNFα | MP6-XT22 | BioLegend | BV605 | 1:100 |
