## Supplemental Methods for "Harnessing Inflammatory Monocytes to Overcome Resistance to Anti-PD-1 Immunotherapy"

### Cell culture.

Tumor cell lines were grown in complete DMEM (Gibco) supplemented with 10% FBS (Gemini Bio-Products) and 1% penicillin/streptomycin (Gibco). All cell lines were periodically verified to be mycoplasma-negative using the LookOut Mycoplasma PCR Detection Kit (Sigma MP0035). Monocytes were cultured in complete RPMI-1640 supplemented with 10% FBS (Gemini Bio-Tools), 1% penicillin/streptomycin (Gibco), 10mM HEPES (Corning), 55 mM 2-mercaptoethanol (Gibco), 1mM sodium pyruvate (Gibco), and 1X MEM NEAA (Corning). Tumor-infiltrating lymphocytes were cultured in complete RPMI-1640 (Gibco) supplemented with 10% FBS (Gemini Bio-Products), 0.5% penicillin/streptomycin (Gibco), 10mM HEPES (Gibco), and 5.5mM 2-mercaptoethanol (Gibco).

### *In vivo* tumor studies.

Tumor length and width were measured using digital calipers and tumor volume was approximated by calculating $(\pi*(\text{length}*\text{width}^2))/6$. Tumor-bearing mice in survival studies were euthanized if tumor volume exceeded 2000mm$^3$, tumor length surpassed 20mm, or body condition score (BCS) was $\leq 2$ or $\geq 4$.

### Flow cytometry.

Cells were blocked in MACS buffer containing TruStain FcX anti-mouse CD16/32 (BioLegend) and 2% normal mouse serum (Jackson ImmunoResearch) for at least 20 minutes at 4°C and then stained for surface proteins using the referenced antibodies (Supplementary Table 2) diluted in MACS buffer for 30 minutes at 4°C. Dead cell exclusion was accomplished using LIVE/DEAD Fixable Viability Dyes (ThermoFisher Scientific). For flow panels including multiple BD brilliant-violet conjugated antibodies, staining master mixes were diluted in BD Horizon Brilliant Stain Buffer (BD). For applications requiring intracellular protein staining, cells were fixed and permeabilized using the eBioscience Intracellular Fixation & Permeabilization Buffer Set (ThermoFisher Scientific). Fixed cells were blocked in 1X eBioscience Permeabilization Buffer supplemented with TruStain FcX anti-mouse CD16/32 (BioLegend) and 2% normal mouse serum (Jackson ImmunoResearch) and subsequently stained with antibodies targeting intracellular proteins diluted in 1X eBioscience Permeabilization Buffer (ThermoFisher Scientific). Stained cells analyzed on an LSRFortessa (BD Biosciences), FACSymphony A3 (BD Biosciences), or Aurora (Cytek Biosciences) flow cytometer. CountBright Absolute Counting Beads (ThermoFisher Scientific) were added to samples pre-analysis to quantify cell numbers. Collected data was analyzed using FlowJo software (v10.10.0, TreeStar). Compensation was manually performed post-hoc using single-color control samples. Gating boundaries were determined and assigned using fluorescence-minus one and/or isotype-matched control antibody samples.

**Mouse single-cell analysis.**

Hashtag-oligo counts were normalized counts using a centered log ratio transformation and implemented in the NormalizeData function in the Seurat package. Due to poor capture of hashtag barcodes per cell, Seurat's default method of hashtag demultiplexing implemented in the function HTODemux poorly resolved hashtag identity when examining hashtag expression levels of singlets, doublets, and negatives. Therefore, the HTODemux method was modified to assign every cell to its most likely hashtag: a hashtag identity was assigned to each cell through *k*-medoid clustering, implemented by the clara function in the R package 'cluster' (v.2.1.4), where *k* is the number of unique hashtag-oligo antibodies used in each sample. After clustering, the density of normalized expression was compared to the hashtag identity to ensure the clustering reflected a reasonable stratification. Samples from the WT CD40 agonist-treated experimental group did not reflect a reasonable stratification post-clustering and, therefore, were manually gated on hashtag-oligo expression to determine hashtag identity.

Normalization, scaling, dimensionality reduction and clustering were performed using Seurat's standard clustering workflow with default parameters of the functions NormalizeData, FindVariableFeatures, ScaleData, RunPCA, and RunUMAP. Where applicable, the first 30 principal components were utilized. Unsupervised clustering was performed with the FindNeighbors and FindClusters functions. Several resolutions of clustering were tested, and a final clustering resolution selected based on the correspondence of clusters with known biological markers. Differential expression analyses between clusters were performed using a Wilcoxon rank-sum test with the FindMarkers function.

Clusters defined from cell type markers were subsetted from the Seurat object and the Seurat workflow was re-run using the same methodology of the full dataset. Putative cell-identity markers were used to identify 69,743 myeloid cells, 25,707 T/NK cells, 1,060 B cells, and 2,827 other cell types. Upon examination of cluster-specific markers, 9,308 and 1,050 doublet cells were removed from the myeloid and T/NK cell subsets, respectively. After the removal of doublet cells, each cluster subset was reprocessed.

All downstream analyses and statistics were performed using functions from tidyverse (v1.3.2) and R (v.4.2.1). Visualizations were created with ggplot2 (v3.4.1) and ComplexHeatmap (v.2.12.1).
